# Circumscribing laser cuts attenuate seizure propagation in a mouse model of focal epilepsy

**DOI:** 10.1101/2021.09.17.460788

**Authors:** Seth Lieberman, Daniel A. Rivera, Ryan Morton, Amrit Hingorani, Teresa L. Southard, Lynn Johnson, Jennifer Reukauf, Ryan E. Radwanski, Mingrui Zhao, Nozomi Nishimura, Oliver Bracko, Theodore H. Schwartz, Chris B. Schaffer

## Abstract

In partial onset epilepsy, seizures arise focally in the brain and often propagate, causing acute behavior changes, chronic cognitive decline, and increased mortality. Patients frequently become refractory to medical management, leaving neurosurgical resection of the seizure focus as a primary treatment, which can cause neurologic deficits. In the cortex, focal seizures are thought to spread through horizontal connections in layers II/III, suggesting that selectively severing these connections could block seizure propagation while preserving normal columnar circuitry and function. We induced focal neocortical epilepsy in mice and used tightly-focused femtosecond-duration laser pulses to create a sub-surface, open-cylinder cut surrounding the seizure focus and severing cortical layers II-IV. We monitored seizure propagation using electrophysiological recordings at the seizure focus and at distant electrodes for 3-8 months. With laser cuts, only 5% of seizures propagated to the distant electrodes, compared to 85% in control animals. Laser cuts also decreased the number of seizures that were initiated, so that the average number of propagated seizures per day decreased from 42 in control mice to 1.5 with laser cuts. Physiologically, these cuts produced a modest decrease in cortical blood flow that recovered within days and, at one month, left a ~20-μm wide scar with increased gliosis and localized inflammatory cell infiltration but minimal collateral damage. When placed over motor cortex, cuts did not cause notable deficits in a skilled reaching task. Femtosecond laser produced sub-surface cuts hold promise as a novel neurosurgical approach for intractable focal cortical epilepsy, as might develop following traumatic brain injury.

**Once sentence summary:** In a mouse model of focal epilepsy, sub-surface laser-produced cuts encircling the seizure focus attenuate propagation without behavioral impairment.

## Introduction

One out of twenty-six people will develop epilepsy at some point in their lifetime and about 50 million people worldwide are currently diagnosed with this disease (*1*–*4*). Epilepsy is also the most common neurologic disorder in dogs (*5*). Epilepsy is defined by recurrent seizures, which are characterized by aberrant, rhythmic, synchronized brain activity that is frequently correlated with involuntary behaviors (*1-4*). Partial onset, or focal, epilepsy is a subclass of the disease characterized by seizure initiation from a consistent, localized region, followed by propagation of the seizure to surrounding brain tissue (*6*). Traumatic brain injury and stroke are the most common causes of focal epilepsy (*7*), but it can also have other etiologies, including cortical neoplasia or cortical dysplasia in children (*7*–*9*). While medical management is preferred, this approach fails in about 45% of focal epilepsy patients either due to initial or developed resistance to pharmacological seizure control (*7*, *10*–*12*). The most effective alternative therapy is to localize the epileptic focus using electroencephalogram (EEG) recordings and then remove the seizure focus through tissue resection (*13*) or damage the tissue at the focus through a laser-based thermal injury (*14*), both of which risk leaving patients with neurologic or behavioral deficits (*12*, *13*, *15*–*17*). The most common location of epileptic foci is the temporal lobe, although partial onset epilepsy can also occur in the occipital, parietal, and frontal lobes (*18*–*24*). For such neocortical focal epilepsies only 32 - 67% of patients became seizure free after resection of the focus, and despite the reduction in seizures a significant percentage of these patients are left with various neurologic deficits including loss of vision, sensory loss, agnosia, dysgraphia or agraphia, acalculia, disturbances in body image, problems with higher order functions, and personality changes, which can be as detrimental to everyday life as the seizures the surgery intended to resolve (*19*–*21*, *25*–*28*).

Recent work has improved the understanding of the underlying physiology behind seizure initiation and propagation. In rodents, acute seizures (e.g. those initiated by focal injection of chemoconvulsants into cortex) have been found to initiate in layer V of the neocortex, spread to overlying neurons in layers II/III, and then propagate away from the seizure focus through these supragranular layers (*29*–*31*). This mechanism has been shown to underlie seizure propagation in multiple animal models of acutely-induced focal seizures (*29*–*34*) and may facilitate the propagation of seizures in humans with focal epilepsy (*29*, *31*, *32*). A less invasive surgical approach, multiple subpial transections, capitalizes on this concept of horizontal seizure propagation through the cortex and uses a metal, hook-shaped instrument to make incisions in the cortex, beneath the surface vasculature, that isolate the seizure focus, with the goal of preventing seizure propagation (*35*). In this procedure, several cuts through the focus are made, by dragging the wire hook bluntly through the brain, which is difficult to control, leading to variable cut angles and significant tissue damage (*36*). Follow up studies found that about two-thirds of patients were seizure-free, but with highly variable outcomes between patients, and up to 30% of patients had significant behavioral and neurologic deficits (*37*–*40*). Although the technique holds great promise, the current execution is difficult to control and not standardized, which has led to it being mostly abandoned at many epilepsy centers. A technique that enabled smaller, sub-surface cuts in the cortex, precisely targeted to specific cortical layers, and resulting in less collateral damage could be used to improve and extend the concept of the multiple subpial transection procedure.

Tightly-focused, infrared wavelength, femtosecond-duration laser pulses enable micrometer-sized cuts to be produced below the surface of the brain without affecting the overlying tissue (*41*, *42*). We previously showed in rats that femtosecond laser cuts in layers II – IV that encircled a chemically-induced, acute seizure focus led to a complete blockage of seizure propagation in 35% of animals and a 36% reduction, on average, in propagation in the remaining animals. These cuts caused minimal damage to the adjacent cortex and preserved the response of neurons inside the encircling cut to a peripheral stimulus (*43*). However, it remains unknown what efficacy such cuts have in chronic focal seizure models that more closely approximate human partial onset epilepsy, and whether these cuts continue to block seizures over time, after the initial injury from the laser cut has resolved.

In this paper, we test the hypothesis that (in part due to limited regeneration in the central nervous system (*44*, *45*) encircling sub-surface laser cuts will lead to long-term reduction of seizure propagation, while the precision of the cuts will prevent significant impacts on normal cortical function. We microinjected iron chloride into layer V of the cortex to induce chronic, focal, neocortical epilepsy in mice and used long-term electrophysiological recordings to compare seizure propagation with and without femtosecond laser cuts that spanned layers II – IV and encircled the seizure focus. We found that laser ablation reduces seizure propagation by 95% while having almost no detectable effect on a skilled forelimb reaching/grasping task. These results suggest that the sub-surface cortical cuts with minimal collateral damage to surrounding tissue produced by tightly-focused femtosecond laser pulses could provide a new approach to treating medically-refractory partial onset epilepsy in patients.

## Results

### Laser cuts in layers II-IV of the cortex encircling an epileptic focus reduced seizure propagation

To test the hypothesis that sub-surface cortical laser cuts circumscribing a seizure focus can attenuate propagation, we produced cuts in craniotomized mice spanning from 550 μm to 70 μm beneath the cortical surface creating an open cylinder with a 1-mm diameter (Movie S1). These cuts were produced using tightly focused, femtosecond-duration laser pulses, which, at sufficient energy, can produce micrometer-scale tissue disruption at the laser focus due to nonlinear absorption of laser energy, without causing significant collateral damage (*42*). We first optimized the cutting speed and depth-dependent laser energy used to produce cuts with a uniform thickness of ~55 μm over the 0.5-mm depth range needed to cut layers II-IV (Fig. S1). These cuts were 85% complete, on average, which was largely dependent on the presence of large surface blood vessels that attenuated the incident laser pulse and blocked ablation from occurring in some locations (Fig. S2). We used these optimized laser parameters — together with manual increases in laser power when focusing directly below a large blood vessel to try to increase the completeness of the cut — for our experiments testing the efficacy of circumscribing cuts for seizure attenuation. After laser cuts were placed in craniotomized mice, FeCl_3_ was microinjected at the center of the cylindrical cut, which led to focally initiated seizures within two weeks. We then placed three electrodes in the bone flap and reimplanted it so that one electrode was at the seizure focus and the other two were at distances of 1 mm and 2 mm from the seizure focus. Beginning about two weeks after surgery, we recorded electrocorticography (ECoG) for 24 hours once a week for 1-33 weeks (most mice 8-12 weeks) (Fig. 1A). We thresholded the ECoG recording from the seizure focus to identify epileptiform events and then manually characterized them as seizures (4,723 events across all mice; e.g. Fig. 1B), polyspikes (41,795 events; e.g. Fig. S3A), or interictal spikes (63,473 events; e.g. Fig. S4A), following standard criteria (*42*). We included animal groups with FeCl_3_ or saline injection and with or without laser cuts, named as follows: epilepsy with laser cuts, epilepsy control (no laser cuts), laser cut control (no epilepsy), and surgical control (no epilepsy and no laser cuts).

**Figure 1.**
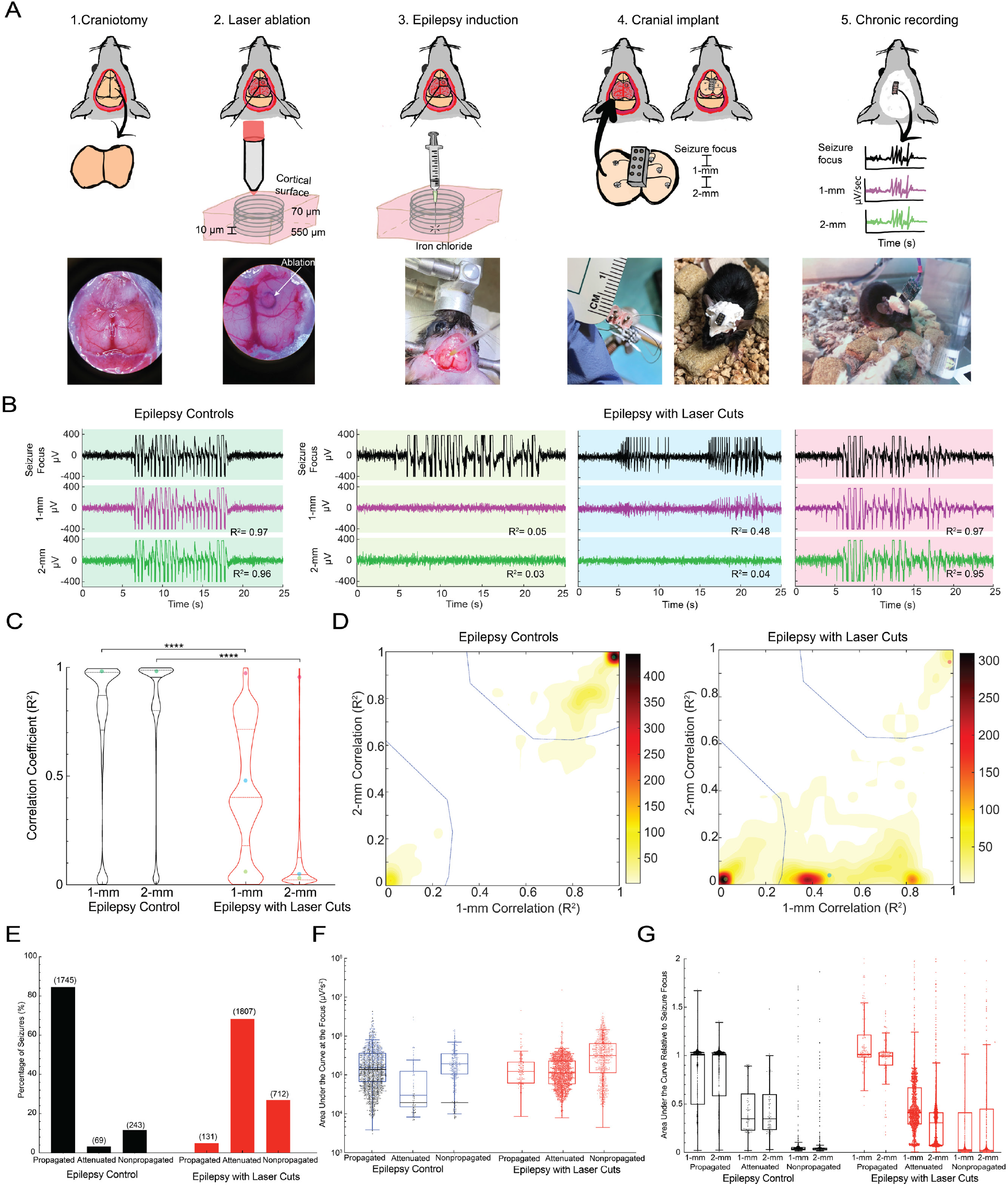
Laser microtransections encircling a chronic seizure focus significantly attenuated or blocked seizure propagation. **(A)** Schematic representation of our surgical procedure. (1) Created a 4 × 10 mm^2^ craniotomy between lambda and bregma. (2). Laser ablation in 1-mm diameter circles at every 10 μm from 550 μm (bottom of layer IV) to 70 μm (top of layer II) beneath the cortical surface. (3) Chronic focal seizures induced by a microinjection of ~350 nL of 100-mM FeCl_3_ in saline into the center of the ablated cylinder. (4) Electrodes implanted in the bone flap at the seizure focus, 1-mm away, and 2-mm away. The bone flap was then reimplanted. (5) Animals were chronically recorded for twenty-four hours once a week for three, or more, months. **(B)** Representative local field potential recordings with correlation coefficients, which indicate how well the recordings at 1-mm and 2-mm distance match the recordings from seizure focus. Background colors are used to indicate where these seizures fall in the plots in C and D (indicated by same color dots). **(C)** Violin plots of all seizure correlation coefficients for epilepsy controls and epilepsy with laser cuts (red) (****P<0.0001, Kolmogorov-Smirnov test). **(D)** 2-D contour maps of the density of seizures as a function of the 1- and 2-mm correlation coefficients, clustered into three groups termed: propagated (top right), attenuated (middle), and non-propagated (bottom left). The representative seizures shown in B include: propagated in a control animal (left), then non-propagated, attenuated, and propagated seizures in a laser cut animal (3 right panels). **(E)** Bar graph indicating the percentage and number of seizures that fell into the three clusters (propagated, attenuated, or non-propagated) for both the epilepsy control and epilepsy with laser cuts. The clustering was statistically different between laser cut and control groups (P<0.0001, Chi-squared test of associations). **(F)** Area under the curve for seizures at the focus comparing the total power of seizures in each cluster between epilepsy with laser cuts (red) and epilepsy control. Dots represent individual seizures (blue dots indicate an outlier animal). **(G)** Area under the curve at distant electrodes comparing clusters (propagated, attenuated, and non-propagated) between epilepsy control and epilepsy with laser cuts (red) (dots represent individual seizures).

Interictal spikes and polyspikes were both regularly recorded in all groups, but at increased incidence in animals with FeCl_3_ injection (Fig. S5). Seizures were nearly absent in the laser cut control and surgical control groups but occurred at an incidence of around 44 +/− 25 and 49 +/− 79 seizures/day in the epilepsy control and epilepsy with laser cut groups, respectively (Fig. S5).

We calculated the correlation coefficient between the ECoG recording at the seizure focus and the ones recorded at distances of 1 and 2 mm (Fig. 1B). In the epilepsy controls, the electrophysiology recording at 1 and 2 mm was highly correlated with the one at the seizure focus, while for epilepsy with laser cuts group, the correlation was reduced and more variable at 1 mm and was reduced even further at 2 mm, causing the distributions of the two groups to be significantly different from one another (P<0.0001, Kolmogorov-Smirnov test) (Fig. 1C). A similar correlation analysis of the propagation of polyspikes (Fig. S3B) and interictal spikes (Fig. S4B) showed significantly reduced correlation coefficients at 1 and 2 mm in the epilepsy with laser cut animals, as compared to the epilepsy controls.

To characterize the propagation of individual epileptiform events, we separately examined seizures (Fig. S6A), polyspikes (Fig. S6B), and interictal spikes (Fig. S6C), combining data from all animals with FeCl_3_ injections. These events were clustered based on the correlation coefficients of the ECoG recording at 1 and 2 mm, relative to the one at the seizure focus, using the Mahalanobis distance-based clustering algorithm (*46*). We first varied the number of clusters and found that using three clusters reliably led to similar group boundaries, more clusters led to groupings that were inconsistent and sometimes nonsensical (e.g. dividing a dense mass of events). With three clusters, events grouped into a compact cluster with high correlation at 1 and 2 mm – a “propagated” event; another cluster with low correlation at 1 and 2 mm – a “non-propagated” event; and into a lower density cluster that tended to have reduced correlation at 1 mm and further reduced correlation at 2 mm – an “attenuated” event (Fig. S6). The laser cuts produced a significant difference in the propagation of epileptic events (Fig. 1D). With the clusters we defined, we found that with laser cuts only 5% of seizures propagated, 68% were attenuated, and 27% did not propagate, as compared to 85% propagated, 3% attenuated, and 12% non-propagated without laser cuts (P<0.0001, Chi-squared test of association) (Fig. 1E). We also found nearly as strong reductions in the propagation of polyspikes (Fig. S3C and D) and interictal spikes (Fig. S4C and D) with laser cuts.

### Seizures blocked by laser cuts were as powerful as propagated ones in controls

To rule out the possibility that attenuated or non-propagated seizures in mice with laser cuts were just less powerful at the seizure focus, we used the ECoG recordings from the seizure focus to calculate several seizure metrics, including maximum band power (over 0 - 50 Hz), seizure duration, and integrated band power over the seizure – termed the area under the curve (AUC). In control mice, attenuated seizures were significantly less powerful than propagated seizures, as expected (Fig. 1F). However, the non-propagated seizures appeared to be equally as powerful as propagated ones. We found that this was dominated by a single control mouse with 87% of the non-propagated seizures (blue data points and median indicators in Fig. 1F). With this mouse excluded, both non-propagated and attenuated seizures had lower AUC, max band power, and duration as compared to propagated seizures in control mice (Fig. 1F, Fig. S6A and B; black median indicators; P<0.0001, one-way ANOVA, Tukey post hoc multiple comparisons correction). In mice with laser cuts, we found no difference in the AUC or duration at the seizure focus between propagated, attenuated, and non-propagated seizures (Fig. 1F, Fig. S7A). The max band power was actually higher in non-propagated seizures, but this was dominated by seizures from a single animal (orange data points and median indicators; Fig. S7B). Critically, there was no difference in the power of propagated, attenuated, or non-propagated seizures in mice with laser cuts when compared with propagated seizures in control mice, indicating that without the laser cuts these seizures would likely have propagated. We also characterized the time of day during the 24-hour recording period that seizures occurred and found no differences between laser cut and control groups or between propagated, attenuated, or non-propagated seizures (Fig. S7C).

The duration, max band power, and AUC of seizures measured at the distant electrodes (and normalized to the measurements from the focus) were each decreased for attenuated and non-propagated seizures, as compared to propagated seizures, for mice both with and without laser cuts (Fig. 1G, Fig. S8A and B; P<0.0001, one-way ANOVA, Tukey post hoc multiple comparisons correction). This suggests clustering seizures based on correlation between focal and distant ECoG recordings distinguished electrophysiologically meaningful propagation, attenuation, and non-propagation. Defining seizure arrival as the time when seizure power exceeded a threshold (average band power over the entire recording session plus one standard deviation), we measured the delay for seizures to propagate from the focus to distant electrodes and found no difference between mice with and without laser cuts (Fig. S8C).

### Additional encircling cuts did not improve blocking efficacy, but revealed that laser cuts reduced seizure initiation

To test whether additional severing of lateral connections increases the percentage of seizures that are blocked, we placed closely (100 μm) (Movie S2) and widely (300 μm) (Movie S3) spaced double cuts around the seizure focus (Fig. 2A). Neither closely nor widely spaced double cuts decreased the percentage of seizures that propagated (Fig. 2B). We did find a trend toward a decreasing number of seizures initiating each day in animals with laser cuts: 44 +/− 25 in epilepsy controls, 49 +/− 79 in mice with single cuts (but with a bimodal distribution), 6 +/− 1 with closely spaced double cuts, and 9 +/− 13 with widely spaced double cuts (Fig 2C). In animals with single laser cuts, there was significant variability in the number of seizures that initiated each day; nonetheless 63% of single cut and 75% of double cut mice had fewer than seven seizures per day, an 85% reduction compared to controls. We did not see similar trends toward decreased numbers of interictal spikes or polyspikes with single cuts or either of the double cut geometries (Fig. S9). The decrease in seizure incidence together with the decrease in seizure propagation led to less than two seizures/day that propagated past the cuts, on average, for all cut geometries (Fig. 2D). These results suggest that making extra cuts does not dramatically improve the attenuation of seizure propagation but may temper seizure initiation.

**Figure 2.**
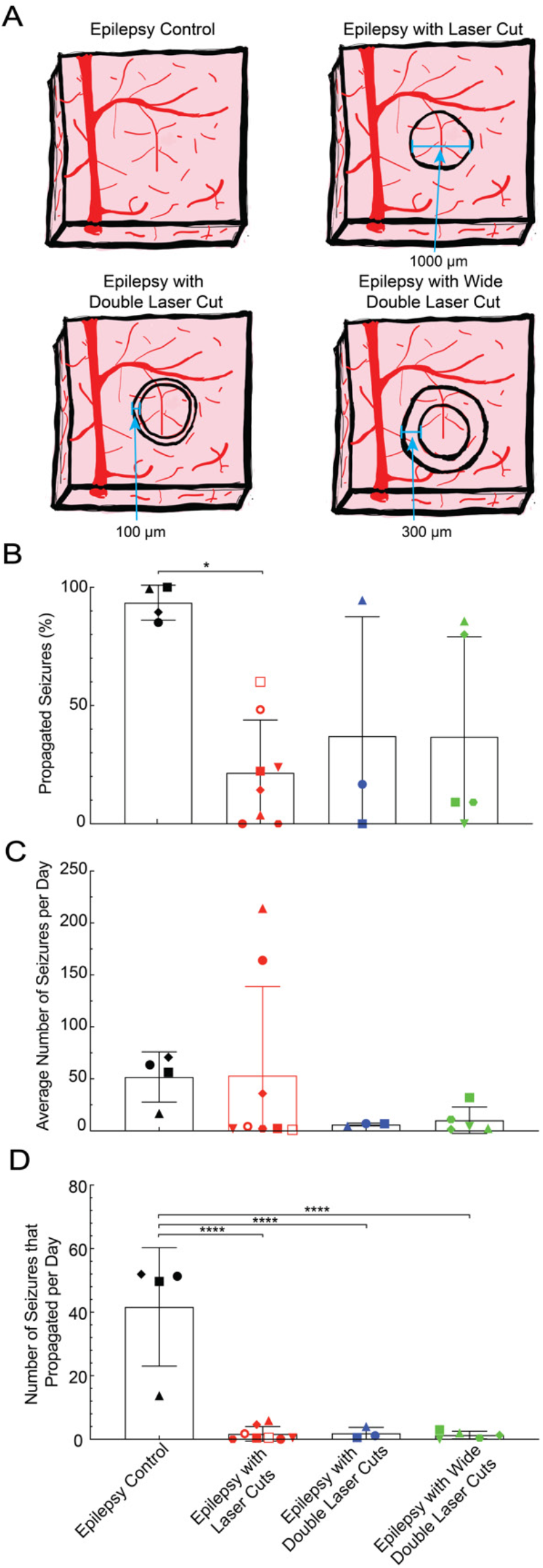
Closely- and widely-spaced double cuts did not further decrease the percentage of propagated seizures but did decrease the number of daily seizures. **(A)** Schematic representation of the different laser cut geometries. **(B)** Percentage of seizures that propagated in epilepsy control, single cut, close double cut, and wide double cut animals (*P=0.01, One-way ANOVA, Tukey post hoc multiple comparisons correction). **(C)** Comparison of the average number of seizures per day between controls and the three types of laser cuts made. **(D)**Average number of seizures per day that propagated in each group (****P<0.0001, One-way ANOVA, Tukey post hoc multiple comparisons correction).

### Seizure blocking efficacy varied across mice, but not over time

The number of seizures and the fraction that propagated varied between recording sessions and between animals (Fig. 3A). Recording sessions from mice without laser cuts tended to have larger numbers of seizures and high propagation, while recording sessions from mice with laser cuts tended to have larger numbers of seizures with reduced propagation, or reduced numbers of seizures and highly variable propagation (Fig. 3A). Averaging across all recording sessions for each mouse, we found that laser cuts reduced seizure propagation from 94% in controls (range: 85% to 100%) to 21% in treated mice (range: 0% to 61%) (p=0.0001, t-test; Fig. 3B). Averaging across all mice, we found that the decreased propagation in laser cut mice did not significantly change out to 12 weeks (Linear mixed model; Fig. 3C). The number of seizures per day trended lower over time, and was similarly variable in mice with and without laser cuts (Fig. 3D), but due to reduced propagation, laser cut mice had, on average, less than two propagated seizures per day, as compared to 42 per day in control mice (P<0.0001, t-test, Fig. 3E).

**Figure 3.**
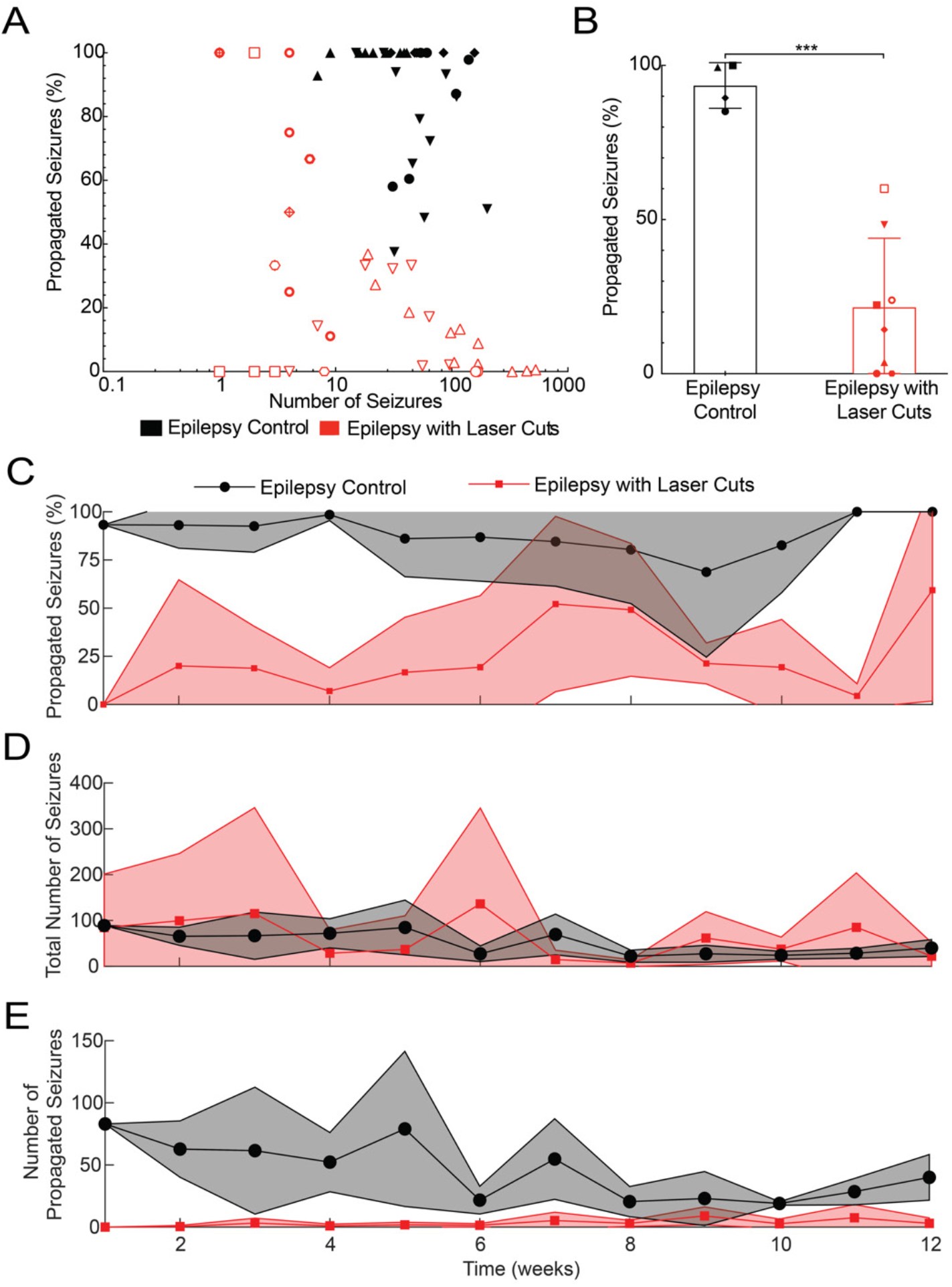
Laser ablation significantly reduced average seizure propagation across all animals and remained effective over time. **(A)** Percentage of seizures that propagated as a function of seizure incidence for both epilepsy controls and epilepsy with laser cuts (red). Each animal is represented by a different shape, with recordings from multiple days for each animal. **(B)** Average seizure propagation for each animal for epilepsy with laser cuts and the epilepsy control groups (***P=0.0001, unpaired t-test). **(C)** Percentage of propagated seizures, **(D)** total number of seizures per day, and **(E)** total number of propagated seizures per day, averaged across all animals, as a function of time for the epilepsy control group and the epilepsy with laser cut group (red).

A few mice had greatly increased numbers of seizures, enabling us to define efficacy of the laser cuts week-by-week for each of these mice over an extended duration. Figure 4 shows seizure propagation data from a mouse with closely spaced double cuts (Fig. 4A and B), one with a single cut (Fig. 4C), and a control (Fig. 4D). In the double cut animal (which had so many seizures it was excluded from the quantitative analysis above to avoid bias), the seizure blocking efficiency improved over several weeks, then regressed back toward the efficacy shortly after the cuts were made where it stabilized from 18 - 24 weeks. For the single cut animal, blocking was highly effective and remained stable over the ~33 weeks. In the control animal, seizure propagation was high and stable over 24 weeks. Interestingly, both the double and single cut mice showed a decreasing number of seizures per day, while seizure incidence was unchanged in the control animal.

**Figure 4.**
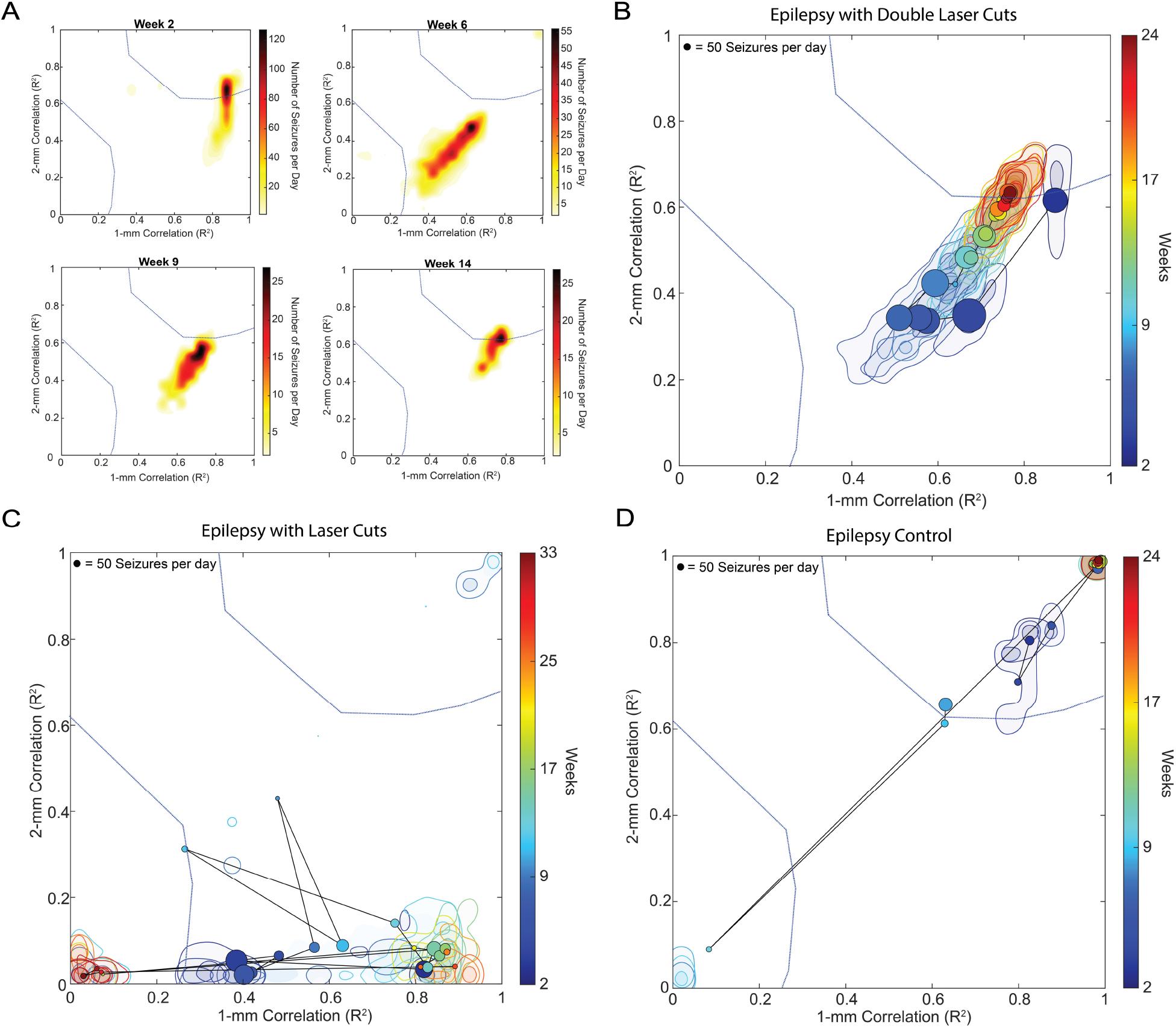
Seizure propagation blocking by laser cuts was stable over time. A closely-spaced double cut, a single cut, and a control animal, each with high incidence of seizures, were followed for an extended period of time to inspect the week-to-week changes in the efficiency of laser cuts in blocking seizures. **(A)** 2-D contour plots of the correlation of recordings of seizures at 1 and 2 mm with the recording from the seizure focus at different times for a mouse with closely-spaced double cuts. **(B-D)** 2-D contour plots of correlation coefficient over time showing the average centroid and the 80^th^ and 33^rd^ percentile contours for the peak density of seizures for each week for mice with a closely-spaced double cut (B), a single cut (C), and no laser cuts (D). The lines connect successive geometric medians of seizure density, with the color of the centroid indicated the recording time and the size of the centroid indicating how many seizures occurred during the 24-hour recording that week (scale in top left corner). To have sufficient data to determine the averages of the correlation coefficients, we excluded data from any recording days that had fewer than 10 seizures.

### Laser cuts cause modest, short-lived blood flow decreases and minor chronic tissue changes

We used three approaches to characterize the physiological and tissue changes that result from the femtosecond laser cuts. First, we took multiphoton images of the tissue before and after producing cuts in mice expressing markers for microglia (CX3CR1-GFP) and neurons (Thy1-YFP) (Fig. 5A). Neurons and blood vessels remained intact at the center of the cut region. Extravasation of blood plasma into the tissue was apparent at the cut border, but nearby blood vessels and neurites remained intact (Movie S1). No blood vessels on the brain surface were disrupted. Tissue was only ablated right at the cut border, where tightly focused femtosecond laser pulses were focused causing multiphoton and avalanche ionization which vaporized tissue (*41*) and led to a small cavitation bubble that displaced tissue (*47*). In the center of the cylindrical cut, microglia and neurite morphologies remained unchanged over hours after the ablation, suggesting minimal acute impacts at this distance from the cut. Overall, there were minimal changes found *in vivo* after laser ablation in regions even tens of micrometers away from the cut locations.

**Figure 5.**
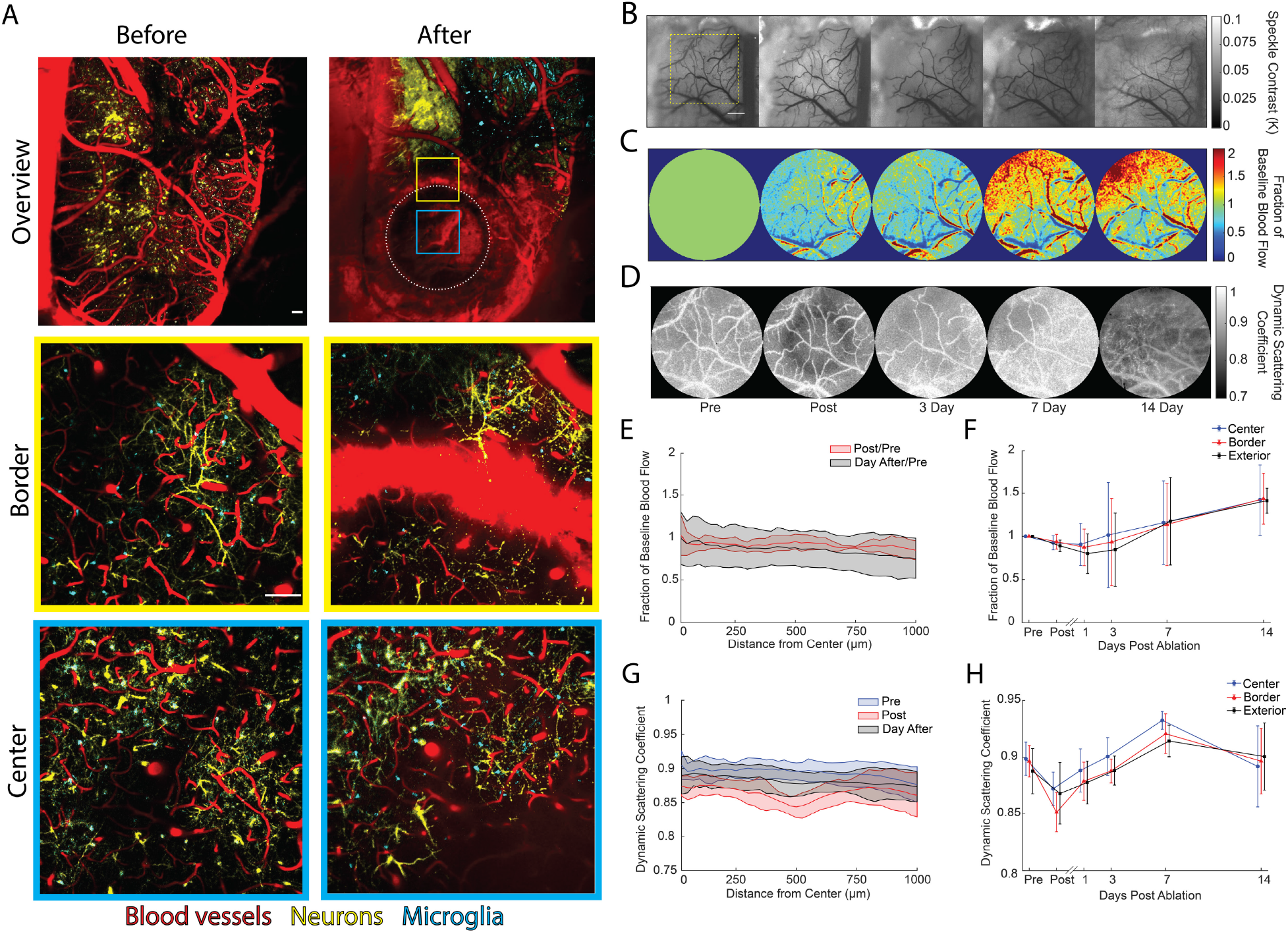
Laser cuts cause minimal collateral damage and only a modest, transient impact on local cortical blood flow. **(A)** Two-photon images showing blood vessels (red), neurons (yellow), and microglia (cyan). Top panel boxes show low magnification images taken before and after producing a 1-mm diameter ablation from 550 μm to 70 μm beneath the surface of the cortex (white dotted circle: ablation, blue box: center, yellow box: border) (scale bar: 100 μm). Middle and bottom panels show zoomed-in views of the border and ablated regions, respectively, before and after ablation (scale bar: 100 μm). **(B)** Multi-exposure speckle contrast imaging captured before, after, and 3, 7, and 14 days after ablation. Images show a single exposure time (25 ms). The ablated region is evident as a faint white ring in the post image (scale bar: 500 μm). Yellow dashed box indicates the field of view for images in panels C and D. **(C)** Heat map showing the fractional change in blood flow over the two weeks recorded. **(D)** Map of the fraction of light scattered by moving scatterers, such as flowing blood cells, over time. The cellular debris from the ablation shows up as a dark ring in the post image. **(E)** Parenchymal blood flow, expressed as a fraction of baseline, as a function of distance from the cut center and measured immediately after and one day after the ablation. **(F)** Fractional changes in baseline parenchymal blood flow for each region (inside the cut, at the border, and outside the cut) as a function of time. **(G)** Fraction of light dynamically scattered as a function of distance from the cut center measured before, immediately after, and the day after ablation. **(H)** Fraction of light dynamically scattered for each region as a function of time.

Second, we used multi-exposure speckle imaging (MESI) to assess the impact of laser ablation on brain tissue perfusion over two weeks post-op in these same mice (Fig. 5B). MESI enables the cortical perfusion (Fig. 5C) and the fraction of optical scatterers that are moving (Fig. 5D) to be extracted from fits to the exposure-time-dependent speckle contrast (*48*–*50*). We first analyzed the impact shortly after the cut as a function of radial distance from the cut center and found only a weak trend toward decreased tissue perfusion with no clear dependence on distance from the cut (Fig. 5E). There was a clear decrease in the fraction of light scattered by moving tissue components near 500 μm from the cut center, at the location of the cut (Fig. 5F). Based on this finding, we examined how perfusion and the degree of dynamic scattering changed over time in regions radially outward relative to the cut center defined as: inside the cut (≤ 450 μm from the center), at the border (450 μm < d ≤ 550 μm), and outside the cut (550 μm < d ≤ 1000 μm). There was a minor blood flow decrease after the cut that recovered to baseline within three days and then increased above baseline flow through the two weeks of observation (Fig. 5G). This increase in flow did not vary strongly across the three regions and was associated with vascular remodeling at the brain surface (evident in speckle contrast images at later times; Fig. 5B). The fraction of dynamically scattered light decreased immediately after the laser cuts, most strongly at the cut border, likely associated with the bleeding and extravasation of red blood cells into the brain tissue at the cut and onto the brain surface (Fig. 5H). The fraction of dynamically scattered light returned to normal within a few days.

Third, we performed histology one month after laser cuts were made to determine the chronic effect of laser cuts on tissue architecture. Coronal sections of the brain, including the area of the laser cut, were examined using histochemical stains and immunohistochemistry. The area of the cut was marked on the cortical surface using surgical ink, and the contralateral cortex was used as a control. Hematoxylin and eosin (H & E) staining revealed linear areas of hypercellularity along the trajectory of the laser cut and in the adjacent leptomeninges, as well as mild disorganization of the neuronal layers in the cortex between the cuts, but no notable disruption of cortical structure (Fig 6A). Tissue fixation has been shown to shrink sections by 25% (*51*), therefore the diameter of the ablated cylinder was found to have shrunk from 1 mm to 0.8 mm and the height of the cut border through layers II - IV shrank from 0.5 mm to 0.4 mm. Despite the shrinkage of tissue, it is still clear when inspecting the cellular organization of the cortex that laser cuts severed layers II/III and cut into the top of layer IV. Acute histology to optimize laser cuts indicated that cuts were 55 μm in thickness immediately after ablation (Fig. S2). Histology one month later showed that cuts left a scar with, on average, 20 μm thickness. At the cut, there was an increase in the density of cells that were immunoreactive for Iba1, a marker of brain microglia and, potentially, invading macrophages (Fig 6B). Prussian blue staining for iron further revealed intracytoplasmic accumulation of hemosiderin in most of these Iba1-positive cells (Fig 6C). Luxol fast blue staining for myelin showed mild myelin loss superficially inside the ablated cylinder (Fig 6D). Immunohistochemistry for olig2 (oligodendrocyte marker) (Fig 6E) and glial fibrillar acidic protein (astrocyte marker) (Fig 6F) showed a very mild increase in these glial cells along the cuts. Cellularity and tissue morphology inside the cuts was similar to the contralateral cortex (Fig 6 A).

**Figure 6.**
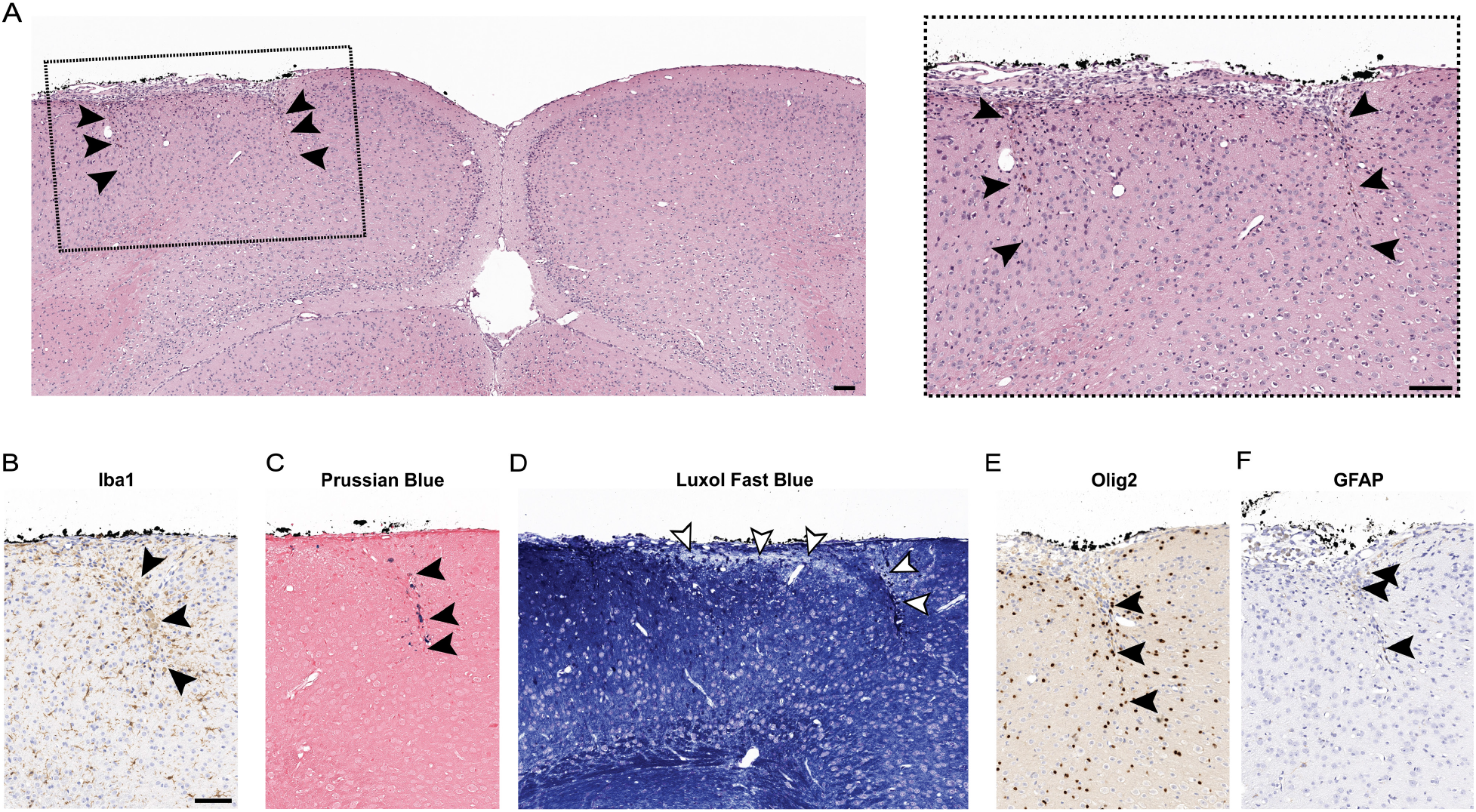
Thirty days after laser cuts the cortical structure was intact and had minute scarring with inflammatory infiltrate, minor gliosis, and mild loss of myelin. **(A)** The left panel shows a light microscope picture of a 5-μm thick coronal section through the cylindrical laser cut, stained with hematoxylin and eosin (H&E), with arrowheads indicating the top, middle, and bottom of the cuts. The dashed box indicates the location for the panel on the right showing a zoomed in image of the laser cuts. Additional panels show histological stains and immunohistological labeling at the location of laser cuts for: **(B)** Iba1 (microglia/macrophages), **(C)** Prussian blue (hemosiderin), **(D)** Luxol fast blue (myelin), **(E)** Olig2 (oligodendrocytes), and **(F)** glial fibrillary acid protein (GFAP; astrocytes) (scale bars: 100 μm).

### Encircling laser cuts in motor cortex did not cause deficits in a complex reaching task

To test for any acute or long-lasting behavioral deficits caused by the laser cuts, we placed cuts in the caudal forelimb area of motor cortex (Fig. S10) and studied performance in a skilled forelimb pellet reaching task (Fig. 7A and B). We compared mice with a regular ablation to mice with sham surgeries and to mice with focal photothrombotic strokes, which were expected to show a deficit (Fig. 7C) (*52*). To assess the impact of a more extensive laser ablation that also severed vertical cortical connections (and approached the invasiveness of resection), we included a group of mice with the same cylindrical laser cut, but with the bottom of the cylinder also severed by laser ablation (Fig. 7C) (Movie S4). We found a significant decrease in the pellet retrieval success (Fig. 7D) and first attempt pellet retrieval success (Fig. 7E) in mice with focal strokes (26% overall success, 11% first attempt success; averaged over the first week) and with severing ablations (21%, 7%), as compared to the sham group (43%, 32%; P<0.0001 for overall success and first attempt success of focal stroke and severing ablation vs sham, one-way ANOVA with Tukey’s multiple comparisons test). This deficit did not show any improvement until three weeks after the surgery, indicating that laser ablation can cause a deficit and that deficit comes from severing both vertical and horizontal connections. In contrast, mice with the standard laser cuts showed only a very modest decrease in pellet retrieval success (39%, 28%; P=0.03 for overall success of regular ablation vs sham) in the early days after surgery (Fig. 7C and D) and had performance matching sham mice at all other time points. There was little variation in the pellet reaching attempts across all groups and time points, except for mice with severing ablations, which made modestly more attempts in the early days after the surgery (Fig. 7F). In conclusion, the laser ablation pattern that effectively blocked seizure propagation did not notably impact performance on a complex forelimb motor task when placed in the relevant cortical region.

**Figure 7.**
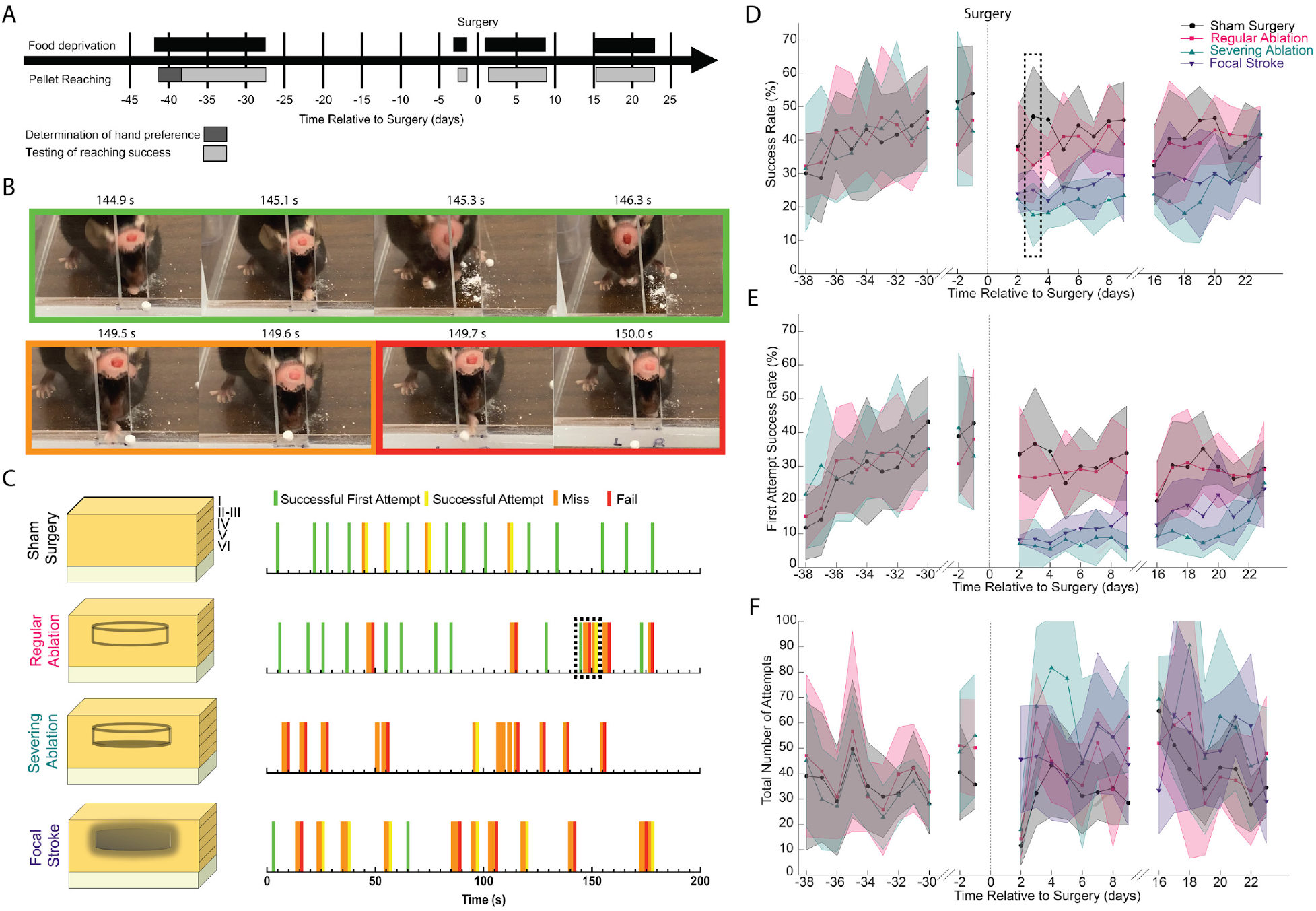
Regular encircling laser ablation targeted to forepaw motor cortex did not reduce long term ability to perform skilled reaching tasks. **(A)** Timeline of when food deprivation and pellet reaching training and testing occurred during the pellet reaching experiment. **(B)** Video snapshots of a first-time successful attempt where an animal grabs and eats the pellet on the first try (green), a miss where an animal reaches and cannot grab the pellet (orange), and a fail where the animal knocks the pellet out of reach (red). **(C)** Schematic representation of a cortical column with the surgical treatment for each of the four experimental groups: sham, regular ablation, a “severing ablation” where the bottom of the cylinder is also laser cut, and a focal photothrombotic stroke. To the right, the panel shows graphical representations of the success of a representative animal’s reaches for a sucrose pellet in the first three minutes of their recording on day 3 post-surgery, for each group. Dashed line box in the regular ablation row indicates the reaches that are shown in panel B. **(D)** Average success rate, **(E)** average first attempt success rate, and **(F)** total number of attempts to grab and eat a pellet during training and after surgery for all four groups (n = 6-7 mice/group). The dashed line box in D indicates the timepoint of the representative data shown in C.

## Discussion

Sub-surface cuts produced by tightly-focused infrared femtosecond laser pulses localized to cortical layers II - IV and encircling a seizure focus were able to block horizontal propagation of focal seizures with high efficacy while minimally affecting normal brain structure or the ability to perform a complex reaching task. With over 100,000 epileptiform events measured in this study, we found laser cuts reduced the propagation of interictal spikes, polyspikes, and especially seizures, for which propagation was reduced from 85% in control animals to 5% in laser cut animals. Seizures that do not propagate, if restricted to a small enough area, may effectively be similar clinically to seizures that never occurred. Moreover, in combination with medication, such cuts might be even more efficacious. There were no differences in seizure power, duration, or other metrics, as measured at the focus, between attenuated and non-propagated seizures in mice with laser cuts and propagated seizures in control mice, indicating that blocked seizures likely would have propagated without the laser cuts. To try and improve the efficacy of this neurosurgical approach we made both close and widely spaced double cuts, but without further notable decreases in seizure propagation. Cuts also tended to reduce the incidence of seizures, with ~70% of all laser cut animals (including both single and double cut) showing an 85% reduction in the average number of daily seizures. This reduction in seizure initiation appeared more robust in animals with double cuts, but this was dominated by two mice with single cuts that had unusually large numbers of seizures, possibly reflecting the variability in seizure incidence previously reported with this model (*53*). Laser cuts also reduced propagated seizures on average by ~97% for all cut geometries when compared with controls (from 42 to 1.5 propagated seizures per day). This reduction in seizure propagation was retained over three months — consistent with the poor regenerative nature of the adult central nervous system (*44*, *45*) and mild glial scarring at the cut, which also may create a barrier to seizure propagation — suggesting that longer-term efficacy may be possible (and which we demonstrated in a few animals that were followed longer). In earlier work, we showed that similar sub-surface laser cuts around an acute seizure focus induced by intracortical injection of a chemoconvulsant reduced both seizure incidence and propagation (*43*, *54*). In comparison, previous studies of the efficacy of the multiple subpial transection procedure in animal models suggested that while the cuts did effectively block kainic-acid induced seizures (*54*) and decreased after-discharges in response to cortical overstimulation (*35*, *55*, *56*), they led to significant neurological deficits.

Unwanted effects on tissue and brain function from these cuts is reduced by the fine cut width, minimal collateral damage, ability to produce sub-surface tissue cuts, and the precise control of the cut location that ablation with tightly-focused femtosecond laser pulses affords (*57*). In our previous work, femtosecond laser cuts were between 80 – 180 μm in width (*43*), which we reduced to ~55 μm while increasing the uniformity and completeness of the cut in this work. These cuts did not injure any blood vessels on the surface of the brain and were completely localized inside the cortex, at the targeted cortical layers. They led to modest decreases in cortical blood flow that recovered in days, consistent with the subsurface nature of the cuts that completely preserved the vasculature on the brain surface. Minor tissue scarring and increased inflammatory cell density was evident at the laser cuts at one month, suggesting the laser cuts do not induce severe chronic pathology. Despite the bleeding from the laser ablation, animals with laser cuts but no FeCl_3_ microinjection did not develop any seizures. The high precision and negligible collateral damage achieved with femtosecond laser ablation is why this tool has been adapted for surgical procedures, including ocular refractive surgery (*57*, *58*), dental cavity removal (*57*, *59*), and removal of bone fragments in orthopedic surgery (under development) (*60*, *61*). We tested the effect of the laser cuts on motor function and found that there were only minor acute deficits in a complex reaching task, which quickly recovered. In contrast, severing vertical connections in conjunction with horizontal connections reduced motor function significantly – comparable to the impact of a focal stroke – and these animals did not quickly recover. These more severe laser cuts are likely closer to the surgical removal of the seizure focus often used clinically, and which can lead to significant neurologic deficits (*20*). Previously, we showed that somatosensory cortex encircled by similar laser cuts still responded to a peripheral stimulus, further supporting the notion of reduced neurological disruption from these cuts (*43*). Thus, the micrometer-sized sub-surface laser cuts that were able to block seizure propagation produced negligible structural damage and nearly absent functional impairment.

Previous work using acute seizure models has shown that focal seizures initiate in the pyramidal cells of layer V of the cortex and then propagate laterally through layers II/III (*29*, *32*–*34*, *62*, *63*). In cortical slices, only cells in layer V were able to initiate the synchronized activity that leads to a seizure (*62*), and only layers II/III were capable of propagating this activity horizontally through the slice (*63*). In both awake and anaesthetized mice, seizures induced in layer V of the cortex relied on layers II/III connections for horizontal propagation (*29*, *32*–*34*). This previous work examining the role of different cortical layers in focal seizure propagation relied on acute seizure models, such as microinjection of 4-aminopyridine. These models reliably generate powerful seizures, but fail to capture the long-term changes in intrinsic neural properties, neural connectivity, or other aspects of the tissue microenvironment that occur during the formation of an epileptic focus (*64*). Such processes could alter the role of different cortical layers in supporting seizure initiation and propagation. The iron chloride model used in this study creates a chronic seizure focus that develops over weeks and models’ aspects of what occurs after a traumatic brain injury (*53*). The significant reduction in propagation of iron chloride induced seizures we found after severing layer II – IV connections is consistent with the idea that seizures in chronic epilepsy also propagate in layer-specific fashion.

About 5% of all epilepsy cases are due to traumatic brain injury (TBI) associated with military service, sports, accidents, or other inciting factors (*65*). A variety of effects secondary to TBI have been proposed to contribute to formation of a seizure focus, including neuroinflammation (*66*), blood-brain barrier permeability (*67*), tissue hypoxia (*68*), changes in neural excitability (*69*, *70*), reactive oxygen mediated damage from cerebral bleeding (*71*, *72*), and metabolic dysfunction (*73*–*75*). The iron chloride microinjection model partially captures the development of a TBI-induced seizure focus by simulating the accumulation of iron from breakdown of hemoglobin left in the tissue after small microhemorrhages from a TBI (*53*, *76*–*79*). The iron accumulation is thought to lead to reactive oxygen species mediated membrane lipid peroxidation preferentially on glial cells, which reduces removal of glutamate from the synaptic cleft, leading to overall hyperexcitability and the eventual development of a seizure focus (*53*, *76*–*79*). Animal models of focal TBI, including the fluid percussion injury and controlled cortical impact models, also lead to focal seizures, but these more complex models normally have fewer than ~1 seizure/day. Future work should investigate the capability of encircling laser cuts to attenuate the seizures that are produced by these models as well.

One remaining question is how to translate this approach into the operating room. With the laser system used in this study we have demonstrated cortical transections as deep as ~1 mm, but this will be inadequate in larger and more complex brains (*42*). The human cortex is not only thicker (1- 4.5 mm) but is also convoluted with gyri and sulci (*80*). This presents two problems: first, the need for increased light penetration to achieve laser ablation at greater depths; and second, the need for a laser delivery system that can traverse the folds of the brain without damaging the surrounding tissue. With longer wavelength laser light, tissue scattering is reduced while linear absorption by water and lipids increases, leading to optimal wavelengths for tissue penetration at the 1.3-μm and 1.7-μm local minima in water absorption (*81*). At these wavelengths, targeted tissue cuts as deep as 5 mm (1.3-μm pulses) and nearly 1 cm (1.7-μm pulses) are theoretically possible (*33*). Using these wavelengths is also likely to decrease the impact of large surface blood vessels on sub-surface ablation, as they did for 2PEF imaging (*82*). Laser technologies that could produce energetic, femtosecond pulses at these wavelengths are becoming increasingly mature (high energy Yb femtosecond laser driving an optical parametric amplifier (*83*) or other nonlinear conversion process (*84*). To enable cortical cuts within the folds of the cortex, a probe that gently penetrates sulci and “side fires” into the cortex could be used. Supporting such a possibility, a flexible, sulci-penetrating needle probe that can enable multiple cortical regions, both on top of gyri and inside sulci, to be reached from a single skull burr hole has recently been demonstrated in cadavers (*85*, *86*). With longer wavelength femtosecond pulses, a sulci-penetrating probe that can focus the light into the tissue and anatomically-registered robotic guidance of probe motion, our procedure could provide a new neurosurgical approach to treating cortical focal epilepsy in patients.

## Materials and Methods

### Study Design

Detailed descriptions of relevant materials and methods are in the Supplementary Materials. All animal experiments were approved by Cornell’s Institutional Animal Care and Use committee. The aim of this study was to determine the effect of encircling micrometer-wide laser cuts that severed lateral connections in layers II-IV of the neocortex on the initiation and propagation of seizures from an induced epileptic focus, as well as to determine the impact of such cuts on the structure and function of surrounding brain tissue. To induce an epileptic focus, we microinjected iron chloride into the cortex of craniotomized mice. Tightly-focused femtosecond laser pulses were then used to produce sub-surface cuts encircling the injection site. Using implanted intracortical electrodes, we measured electrophysiological ECoG recordings from the seizure focus and at electrodes placed 1 and 2 mm away for 12-33 weeks and quantified seizure propagation by examining the correlation of the distant recordings with the seizure focus. We examined four groups: epilepsy controls (n = 4) (induced with seizures but did not have laser cuts), epilepsy with laser cuts (n = 8) (induced with seizures and had laser cuts), laser cut controls (n = 3) (given a sham intracortical injection, but had laser cuts), and surgical controls (n = 3) (given a sham intracortical injection and had no laser cuts). We further tested the effect of adding concentric laser cuts with “close” (n = 4) and “wide” (n = 5) spacing in mice with epilepsy. To assess the effects of laser cuts on the structure and function of the cortex, we used several approaches. We imaged mice (n = 5) with fluorescently labeled neurons and microglia using multiphoton microscopy before and after ablation to see the acute effect on tissue architecture. We followed cortical blood flow changes in the same mice over two weeks after ablation using multi-exposure laser speckle imaging. In mice that had laser cuts for about a month, we histologically examined impacts on tissue architecture and cellular composition near the cuts. Lastly, we examined the impact of cuts placed in motor cortex on success in a skilled forelimb reaching/grasping task. Mice were broken into three groups: sham surgery (n = 6) (just a craniotomy), regular ablation (n = 6) (laser cuts were placed in caudal forelimb motor cortex), and severing ablation (n = 7) (laser cuts in the same place, but with the addition of a full ablated layer at the bottom of the cylinder, thereby severing vertical cortical connections). In the task, mice learned to grasp a sugar pellet through a narrow opening, and we assessed changes in their success after treatment. Sham surgery mice repeated the experiment after being given a photothrombotic stroke of similar size to the laser ablated region, as a positive control. For the seizure propagation and grasping task experiments, group sizes were determined from power analysis based on results from pilot experiments (see Supplementary Methods).

## Supporting information

Supplementary Move 2

## Supplementary Materials

Detailed Materials and Methods

Figs. S1 – S11

Movies S1 – S4

## Acknowledgments

We gratefully acknowledge the extra care taken by the animal facility staff with the mice, most with complex implants and surgical procedures, used in this study.

## Funding

This work received support from the National Institutes of Neurological Disorders and Stroke R21 NS078644 (CBS and THS), the National Institutes of Mental Health R43 MH119880 (CBS), the Clinical Translational Science Center National Center for Advancing Translational Sciences UL1 RR024996 pilot grant (MZ and CBS), the Cornell University Ithaca-WCMC seed grant (MZ and CBS), and the Daedalus Fund for Innovation (THS, CBS, and MZ).

## Author contributions

Conceptualization: SL, THS, CBS

Methodology: SL, DAR, RER, MZ, NN, OB, THS, CBS

Investigation: SL, DAR, RM, AH, JR, TLS, OB

Software: DAR

Visualization: SL, DAR, CBS

Formal analysis: SL, DAR, TLS, LJ, NN, CBS

Funding acquisition: MZ, THS, CBS

Project administration: THS, CBS

Writing – original draft: SL, DAR

Writing – review & editing: SL, DAR, RM, AH, TLS, LJ, JR, RER, MZ, NN, OB, THS, CBS

## Competing interests

None

## Data and materials availability

The data underlying each figure or result will be uploaded to Cornell’s eCommons and will be publicly available.

## Notes

### Competing Interest Statement

The authors have declared no competing interest.

