## Supplementary Materials for "Circumscribing laser cuts attenuate seizure propagation in a mouse model of focal epilepsy"

**Detailed Materials and Methods**

1. *Surgery and electrophysiological recording for evaluation of long-term impact of laser ablation on seizure propagation*

All animal procedures were approved by the Cornell Institutional Animal Care and Use Committee (protocol number 2015-0029) and were conducted following NIH guidelines. These experiments required a staged animal preparation procedure. First, access to the neocortex was obtained by performing a craniotomy (Fig. 1A1). Next, we used two-photon microscopy to map the cortex and guide the production of a 1-mm diameter cylindrically-shaped sub-surface cut through targeted cortical layers using tightly-focused femtosecond duration laser pulses, in order to sever lateral neural connections (Fig. 1A2). We then microinjected iron chloride into the cortex at the center of the cut region, which drives the formation of a seizure focus within 1-2 weeks (Fig. 1A3). Finally, electrodes were implanted over the craniotomy to record from the seizure focus, and at two sequential linear distant locations that sit outside the cylindrical cut (Fig. 1A4). Mice with only iron chloride injections, with only laser cuts, or only the electrodes served as controls. We then took freely behaving electrophysiological recordings for 24 hours, weekly for 1-33 weeks (>8 weeks for 75% of mice), and quantified the incidence and propagation characteristics of epileptiform events. The sections below provide detailed descriptions of these experimental steps.

**1.1. Animal groups and preparation for surgery**

All animals were purchased from Jackson Laboratory (strain: 000664-B6, Jackson Laboratory) or locally-bred C57BL/6J wildtype mice, ranging from 18 - 40 g in weight, between 2 and 12 months in age, and split between male and female animals in each experimental and control group. Animals were split into four groups with three to eight animals per group: an experimental group which received laser ablation, iron chloride injection, and electrode implantation (n = 8, 5 male and 3 female) ; an epilepsy control group which received iron chloride injection, electrode implantation but no laser ablation (n = 4, 3 male and 1 female); a laser cut control group which received laser ablation, saline injection, and electrode implantation (n = 3, 1 male and 2 female); and a surgical control group which received saline injection, electrode implantation, but no laser ablation (n = 3, 2 male and 1 female). Group sizes were determined based on preliminary data from pilot experiments that showed epilepsy control animals would have around 25 +/- 10 seizures a day. A power analysis was run with GPower to determine statistical significance at a p = 0.05 level for a 10% decrease in seizure propagation frequency, suggesting about 700 seizures were needed. We achieved this with a minimum of 4 animals per group with at least 8 weekly recording sessions per animal (~200 seizures per animal). Laser cut controls and surgical controls were deemed sufficient when the first three animals showed essentially no seizures. Data collection was stopped when 4 animals in both the epilepsy control and epilepsy with laser cuts had made it through at least 8 -12 weeks of recordings.

All surgeries were performed at the same time of day using the same tools and in the same location within the laboratory. Animals were induced using 3-4% isoflurane in 100% oxygen through a nose cone and then maintained at 2-2.5% isoflurane with minor adjustments to maintain a respiration rate of ~1 Hz. Every hour during the surgical procedure the animals were given subcutaneous injections of atropine (0.005 mg/100 g mouse weight; 54925-063-10, Med Pharmex) to help suppress excess lung secretions, and 5% glucose (1 mL/100 g mouse weight) in saline to prevent hypovolemia. The animal's temperature was maintained at 37ºC using a feedback-controlled heating pad (40-90-8D DC, FHC). A large (~4x10 mm$^2$) craniotomy was opened using a dental drill to expose the cortex. The removed bone piece was maintained in sterile saline for later reimplantation with inserted recording electrodes. The animals in the epilepsy-only and surgical control groups were immediately prepared for intracortical injection by applying saline to the exposed brain and preparing the Nanoinject II (Drummond). The animals in epilepsy with laser cut and laser control groups had the craniotomy temporarily closed with a 13-mm diameter circular cover glass, with saline underneath, that was lightly adhered with cyanoacrylate adhesive (Loctite 420) at the rostral and caudal edges. The rim of the craniotomy was then sealed using Kwik-Sil (World Precision Instruments) and the animals were moved to the microscope for imaging and ablation.

#### 1.2. Imaging and ablation procedure

For the ablation procedure, mice were first retro-orbitally injected with Texas Red dextran (40 µl, 2.5% w/v, molecular weight = 70,000 kDa, Thermo Fisher Scientific) in saline to label the vasculature. Mice were then moved to a custom built two-photon excited fluorescence (2PEF) microscope that utilized a 1,030-nm femtosecond laser for excitation (Satsuma, Amplitude Systemes). Texas Red emission was collected on a photomultiplier tube (PMT) through a 630/92 nm (center wavelength/bandwidth) emission filter. Laser scanning and data acquisition was controlled using ScanImage software (*87*). We collected low magnification images of the brain surface vasculature using a 0.28 NA, 4X objective (Olympus). This ~2.5 x 2.5 $mm^2$ vascular map enabled us to reliably identify the same brain regions under both the 2PEF and the surgical dissection microscope. We then switched to a higher numerical aperture 0.95 NA, 20x objective (Olympus) for tissue ablation. Ablation utilized 800-nm wavelength, ~50-fs duration pulses from a 1-kHz repetition rate Ti:Sapphire laser amplifier (Legend, Coherent), routed into the 2PEF microscope as previously described (*42, 43*). Briefly, the beam from the amplifier was passed through a λ/2 waveplate, with computer-controlled rotation, and then a polarizing beamsplitter cube in order to adjust laser power. We used an 875-nm long-pass dichroic mirror (FF875-Di01, AVR Optics), placed between the scan and tube lenses, to route the ablation beam into the microscope. The beam first passed through a -150-mm focal length lens that, in combination with the 300-mm focal length tube lens, expanded the ablation beam to fill the back aperture of the objective. The mouse was mounted on a computer-controlled 3D translation stage (XPS, Newport). Custom MATLAB software controlled stage movement, shutters for both the amplified beam and the 2PEF beam, as well as laser powers. We identified ~1-mm diameter regions for ablation, being careful to obtain regions that avoided large overlying blood vessels that could disrupt the ablation beam. All ablated regions were located between 0.5 mm and 4 mm lateral from and left relative to the central sinus and less than 1 mm caudal from bregma, areas encompassing motor cortex and somatosensory barrel cortex (*88*). We created an open top and bottom cylinder with ablated walls by producing concentric circular cuts from ~550 µm to ~70 µm beneath the cortical surface (layers II-IV of neocortex) (*89*), with 10 µm steps. Laser energy was decreased exponentially with decreasing depth (*42*), starting at ~30 µJ at 550 µm and ending at ~4 µJ at 70 µm below the cortical surface to achieve sub-surface cuts to the tissue with uniform width (Fig S1D). We moved the stage at 700 µm/s in a 1-mm diameter circle while irradiating and blocked the amplifier beam when transitioning between layers. In some animals a second layer of cuts were made simultaneously at each layer creating a larger diameter circle outside the 1-mm diameter circle (closely-spaced: 1.2 mm diameter (n = 4, 2 male and 2 female); widely-spaced: 1.6 mm diameter (n = 5, 2 male and 3 female). We also monitored, in real time, the success of the laser ablation by visualizing the rupture of cortical capillaries that fell in the ablation path using 2PEF imaging (Fig S2A; Movie S1). In this and previous work, we found that the presence of large surface vessels interferes with ablation efficiency and therefore we manually increased the ablation power to maintain successful ablation in such conditions. Mice were removed immediately after ablation and returned to surgery.

#### 1.3. Intracortical injection:

Animals in the experimental and laser control groups had the cover glass over the craniotomy carefully removed. All animals had saline-soaked gel foam (Ethicon: 1969) applied to the brain to keep it moist. We pulled glass pipettes and then broke the tips to create micropipettes with ~10-µm openings, which were filled with mineral oil and then backfilled with either 100 mM $FeCl_3$ in saline or just sterile saline. The micropipette was connected to a Nanoinject II (Drummond) mounted on a micromanipulator and was lowered to a depth of ~550 µm beneath the cortical surface (~layer V of the cortex) (*89*). For animals in the experimental and laser cut control groups the pipette was directed to the center of the ablated cylinder, while for animals in the surgical control and epilepsy control groups the pipette was placed at anatomically equivalent locations. We injected ~350 nL (rate: 46 nL/s) of either $FeCl_3$ for the experimental group and epilepsy-only control or saline for the laser cut control and surgical control (*53*).

#### 1.4. Electrode implantation:

The skull fragments removed during the craniotomy were stored in saline until electrode implantation. The bone was placed in a sterile field and burr holes were drilled to hold electrodes. We placed three sequential burr holes in the bone fragment so that, when it was re-implanted, the electrodes were located over the injection site and at distances of 1 and 2 mm away. Two additional burr holes were created over the contralateral hemisphere for a common reference and a ground. Electrodes (0.10" screws with wire leads, Pinnacle Technology) were implanted into the burr holes and soldered to an 8-pin head mount (Pinnacle Technology). The wire leads were shortened and soldered to bring the head mount as close as possible to the bone fragment. Electrodes, head mount, and bone fragment are then together lowered utilizing a micromanipulator for precision back over the brain being careful to align the first electrode right over the injection site. The implant was then sealed using Kwik-Sil (World Precision Instruments) and then cemented using dental cement (C&B metabond) that was mounded up around the implant to secure it to the head for the next several months. Animals then recovered for 1-2 weeks while the epilepsy model developed, and seizures began occurring.

#### 1.5. Electrophysiology:

Electrophysiological recordings were taken using a three-channel tethered system with a sampling rate of 400 Hz from the intracranial surface electrodes, together with simultaneous video recording (Pinnacle Technology). Electrophysiological and video recordings in every animal were taken for twenty-four consecutive hours once a week every week for between one and 33 weeks. All recordings were started around 8 am and lasted for a full 24-hour cycle. A red-light lamp was used to illuminate animals at night for the video recording.

Mice were excluded from the study if seizures did not develop after intracortical injection of iron chloride or if other aspects of the implantation surgery failed (23 mice from epilepsy control group and 10 mice from epilepsy with laser cuts group). This variability in epilepsy incidence after iron chloride is larger than reported previously with this model (*53*). Animals were also excluded if they did not survive long enough for at least one recording session (2 mice from surgery control and laser cut control groups, each). Data from individual recordings was excluded if it was too noisy. The electrode frequency spectrum was computed using a Fourier transform and averaged across 3-s time windows over the entire recording duration. Electrode channels were excluded when 60 ± 0.67 Hz frequencies constituted greater than 10% percent of the total frequency spectrum. Noisy channels appeared over time and were consistently noisy after appearing. Only one closely-spaced double cut animal was removed entirely from the study as an outlier because the mouse showed a significantly higher incidence of seizures and would have skewed the overall data if included. Seizure propagation in this mouse was analyzed separately (Fig. 4A and B)

#### 1.6. Seizure Event Detection and Classification:

We used semi-automated approaches to identify episodes of epileptiform activity in the ~7,500 hours of electrophysiological recordings acquired in this study. To locate potential epileptiform events, a 0.25-s long root-mean-square (RMS) moving window filter was applied to the signal from the electrode at the seizure focus. Peak RMS values exceeding 60 μV were identified. Sequential peaks that were within 1 s of each other were combined into a single event. Because normal activity sometimes exceeded this threshold, we discarded events that did not contain any spiking in the unfiltered signal from the seizure focus that did not exceed 250 μV. A researcher blinded to the treatment of the animal then manually classified each of these events, looking only at the recording from the seizure focus, as an interictal spike (single spike), polyspikes (spiking shorter than 3-s duration), a seizure (spiking longer than 3-s duration), or a false positive (e.g. signal due to motion artifact) (*90*). Simultaneous video was recorded and used to help identify false positives by identifying scratching epochs or the animal transiently shorting electrodes by hitting the enclosure. We then calculated the cross correlation of the electrophysiological recordings taken at distances of 1 and 2 mm with the recording from the seizure focus, called R1 and R2, respectively (allowing for small temporal shifts between the recordings and finding the shift that yields the maximal cross correlation value).

We combined all seizures across all experimental groups and clustered them using the Mahalanobis distance-based algorithm based on these correlation values (*46*). This was repeated for polyspikes and interictal spikes. For the seizures, we found two clear and compact groupings, which we termed propagated seizures (approximately: R1 ~ R2 > 0.65) and non-propagated seizures (R1 ~ R2 < 0.25). For the remaining events, we tried different numbers of clusters, but found that using more than one additional cluster led to inconsistent clustering and groupings that seemed unintuitive. We thus used three clusters total and termed the third cluster attenuated seizures (R1 < 0.85, R2 < 0.25). We similarly arrived at three groupings, with quite similar boundaries, for polyspikes and interictal spikes.

Using these recordings, we also characterized seizure duration and band power, as well as the delay for the seizure to propagate from the focus to distant electrodes. We first determined a threshold power, above which activity was determined to be epileptic. We calculated the signal band power (RMS signal strength averaged over a fixed frequency range and time interval) between 0 and 50 Hz using sequential 1-s intervals across data from the entire recording session and set the threshold for each channel to be the average band power plus one standard deviation. We then calculated band power for each seizure event, again using 1-s interval and 0 to 50 Hz bandwidth, but now with a 25-ms step size through the data (to improve temporal resolution). The channel-by-channel thresholds were then applied to this data to determine the beginning and end of each seizure event. From this we determined the seizure duration, as well as any delay in seizure onset between the focus and distant electrodes. We further calculated the area under the curve (AUC) as the sum of the band power over the duration of the seizure, and the maximum band power during the event.

In these electrophysiological recordings, more powerful seizures saturated the detection electronics. This was deemed better than having too little amplification and missing the propagation of weaker seizures. To determine the effect of signal saturation on the calculated correlation coefficient, we took signals from 11 seizures with no saturated voltages (Seizure) as well as normal ECoG signals from a distant electrode within 5 min. of the event within the same recording session as the non-propagated signal (Distant) (Supplementary Fig. 11). Mixed signals, representing propagated seizures, of different seizure amplitude (but similar noise) were created by linearly summing the seizure signal with the non-propagated signal as Mixed = (n)Seizure + (1-n)Distant, where $0 \leq n \leq 1$, while saturation was controlled by determining a threshold for which the seizure signal reached a set percentage of samples at the positive or negative threshold bounds and applying the threshold to both the Seizure (representing the focus) and Mixed (representing the propagated) signals. We calculated the correlation coefficients between the Seizure and Mixed signals while varying n (e.g. the real amplitude of the propagated seizure) and the degree of saturation in the seizure signal for each trial. We averaged trials from the 11 identified seizures together, and determined the resulting correlation coefficient underestimation based on the 0% saturation condition.

### 2. *Chronic imaging for determining the long-term effects of ablation on tissue architecture and perfusion*

#### 2.1. Surgical procedure:

Transgenic animals (B6CgTg(ThyYFPH) (The Jackson Laboratory stock number: 003782) x B6.129P2(*91*)-Cx3cr1tm1Litt/J (The Jackson Laboratory stock number: 005582)) (*91*) were used to label neurons (Thy1-YFPH) and microglia (CX3CR1-GFP) (n = 9, 6 males and 3 females). Animals were at least 2 months old, between 18-40 g. Animals received a 4-mm permanent cranial window between the suture lines lambda and bregma as described above with a 6-mm diameter circular cover glass fitted over the site and adhered using superglue and dental cement. A small layer of fibrosis tissue and scarring did occur over the craniotomy in four of the nine mice and the layer was removed and new glass cover slip fitted to the head in order to maintain image clarity.

#### 2.2. Chronic multiphoton imaging:

Animals were chronically imaged using both 2PEF and laser speckle contrast imaging microscopes before and after ablation to determine the effect of ablation on tissue architecture and cortical blood flow. Mice were injected with Texas Red dextran (40 µl, 2.5%, molecular weight = 70,000 kDa, Thermo Fisher Scientific) and imaged with a Ti:Sapphire femtosecond laser (Mira, Coherent) set to 880 nm for immediately before and after ablation. The GFP, YFP, and Texas Red fluorescence was collected with a 735-nm long-pass dichroic, separated by secondary and tertiary long-pass dichroics at 520 nm and 593 nm, and detected through bandpass filters with center wavelength/bandwidth of 494/41, 550/49, and 630/92 nm, respectively. Three-dimensional overview imaging stacks were taken of the ablated region before and after ablation using a 4x 0.28 NA objective (XLFLuor4x/340, Olympus) to create ~2.5 x 2.5 $mm^2$ overview. More detailed ~0.5 x 0.5 mm sized 3D image stacks were taken at the center and at the border of the ablated region using a 20x 0.95 NA objective (XLUMPLFLN20XW, Olympus). The ablation procedure remained the same as described above. As fluorophores labeled distinct structures with no anticipated overlap, multiphoton images were spectrally unmixed using the LUMoS unmixing method (*92*) in which image data is clustered based on the number of fluorophores used, then the pixel values are scaled based on the original image intensities.

#### 2.3. Multi-exposure laser speckle imaging:

We used multi-exposure speckle imaging (MESI) (*49*) to assess the impact of the laser cuts on tissue perfusion over two weeks following laser ablation. For imaging, a beam from a 785-nm laser diode (LD785-SEV300, ThorLabs) was variably diffracted through an acousto-optic modulator (AOMO 3100-125, Gooch & Housego), then expanded for widefield imaging on the mouse cortex. Thirty laser speckle images taken at fifteen different exposure times between 50 µs and 80 ms were acquired through a 4X 0.28NA objective (XLFluor, Olympus) onto a CMOS camera (acA2040-90umNIR, Basler). A linear polarizer and IR bandpass filter were placed in front of the CMOS sensor to increase contrast and reduce ambient light levels at the longer exposure times, respectively. A DAQ board (USB-6001, National Instruments) was used to adjust the laser power diffracted through the modulator to maintain a constant image intensity across all exposure times, while a microcontroller gated and synchronized the modulator output with the camera exposure. Dark images with no laser illumination were interleaved with these images to quantify camera noise. Speckle contrast was calculated as the standard deviation divided by the mean ($K = \frac{\sigma_s}{\langle I \rangle}$) in a moving 7x7 pixel window, and using the camera noise reduction approach from Wang, et al. (*93*). Following the work of Postnov, et al. (*94*), pixel contrast values at each exposure time were fit to

$$\begin{aligned} K^2 = \; & (1 - D_{MU})\left(\beta\rho^2 \frac{e^{-2x} - 1 + 2x}{2x^2} + \beta\rho(1-\rho)\frac{e^{-x} - 1 + x}{x^2}\right) + \\ & D_{MU}\left(\beta\rho^2 \frac{e^{-2\sqrt{x}}(4x + 6\sqrt{x} + 3) + 2x - 3}{2x^2} + \beta\rho(1-\rho)\frac{e^{-\sqrt{x}}(2x + 6\sqrt{x} + 6) + x - 6}{x^2}\right) + \\ & \beta(1-\rho)^2 + \nu \end{aligned} \qquad \text{Eq.1}$$

where β is the normalization factor to account for mismatch between the pixel size and the speckle size on the camera, ρ is the fraction of scattered light that is dynamically scattered, and ν is the instrumentation noise. $D_{MU}$ indicates whether the scattering motion by blood cells is ordered like in vessels ($D_{MU}$ = 0) or disordered like in capillary-dense parenchymal regions ($D_{MU}$ = 1). $x = T/\tau_c$, where $T$ is the exposure time and $\tau_c$ is the speckle correlation time which is inversely proportional to the blood flow speed or perfusion.

Measurements were taken prior to the laser cuts, immediately after, then at 1, 3, 7, and 14 days after ablation. We used 50-µs and 80-ms exposure time speckle contrast images to locate the border of the laser cut and thus to define the center, where the increase in non-moving red blood cells in the tissue at the cut border provided signal changes that were readily apparent. We calculated parenchymal tissue perfusion and the fraction of light scattered by dynamic (e.g. moving) scatterers for 25-µm wide concentric rings up to 1 mm out from the center of the cut. In order to average over only parenchymal tissue, a vessel mask was created using the 80-ms exposure speckle contrast images and large vessels were excluded to reduce the probability of measurements from large vessels skewing the results. Two mice were excluded from speckle imaging due to being "over ablated," where excessive laser power was inadvertently used and much wider laser ablation with increased intraparenchymal bleeding was observed, and two more were excluded due to instrumentation issues with the imaging setup during baseline recording.

3. *Acute in vivo imaging and histology to determine optimal cut speed and cut completeness*

**3.1. Surgical procedures:**

All animals were C57BL/6J wildtype mice (18-40 g) bred in our colony and at least 2 months old (n=30). Animals were fitted with a cranial window and laser ablations were performed in the cortex using the same 2PEF imaging and ablation setup described above. Immediately after ablations occurred, animals were transcardially perfused using 4% paraformaldehyde (PFA) and brains were removed for histologic analysis.

**3.2. Ablation procedure for optimal cut speed:**

To find an optimal cutting speed with the 1-kHz laser pulse train, the sample stage was programmed to move the laser focus in single lines of 1-mm length with rostral-caudal orientation at specified depths in the cortex, using the depth-dependent energy found to produce about 50-μm wide cuts in our previous work (*42*). Each of 25 mice received ablations at three different depths in four separate columns, two on each hemisphere of the parietal cortex. Cut speeds varied across 100, 300, 500, 700, or 900 μm/s, and cuts were made at depths of 600, 400, and 200 μm below the surface of the brain. We volumetrically imaged (Texas red dextran was intravenously injected, as above) a subset of the cut lines using an Olympus 20x 0.95NA objective to determine *in vivo* approximation of cut width to compare with histologic analysis.

**3.3. Ablation procedure for cut completeness:**

The ablation laser and 2PEF imaging were set up, five animals were retro-orbitally injected with Texas red dextran, and a vascular map of the craniotomy was obtained using in vivo imaging, as described above. Circular laser cuts were created on both hemispheres having a 1-mm diameter from 550 μm to 70 μm below the surface of the brain. Animals were imaged after ablation with the 0.28 NA 4x Olympus and Olympus 20x 0.95NA objectives to create volumetric stacks (~2.5 x 2.5mm$^2$; ~0.5 x 0.5 mm$^2$) for in vivo visualization of acute cuts.

**3.4. Histologic and image analysis:**

After ablation animals were perfused using 4% PFA in PBS and brains were extracted from the skull before being stored in 4% PFA for 24 hours. Brains were then switched to a solution of 30% sucrose in PBS for another 24 hours. Next, brains were frozen in optimal cutting temperature (OCT) compound (Fisher Healthcare) and sectioned in 25-μm thick sections using a microtome cryostat (Microm HM 550, Thermo Fisher Scientific). Animals with laser cuts from the cut speed experiments were sliced into coronal sections and animals with circular cuts to determine cut completeness were sectioned using horizontal sections to show the entire circle. Sections were mounted on slides and then stained using 3,3'-Diaminobenzidine (DAB) to highlight red blood cells, in order to show the cut regions more clearly. Images were taken utilizing a one-photon microscope (Zeiss Axio Examiner, with a QICam camera) and image analysis was conducted in ImageJ.

Images of line cuts in the coronal sections showed the cross section, allowing measurement of the cut width to determine cutting speeds that minimized collateral damage and verifying the appropriate depth-dependent laser energy. Images of circular cuts in the horizontal sections showed the 1-mm diameter circle where ablation was attempted, allowing the completeness to be measured by manually identifying gaps in the cut. In order to account for misalignment of the cutting plane with the cylinder axis when slicing, the first and last 100 μm of tissue containing circular cuts were excluded, so the cut completeness assay focused on the internal 300 μm.

4. *Pellet reaching task to assess the effect of laser ablation on fine motor function of mice*

**4.1. Animals**

All animals (n = 24) were C57BL/6J mice (strain: 000664-B6, Jackson Laboratory) (12 female, 12 male) between 4 and 5 months of age. Mice were housed in cages with Alpha-Dri bedding (to avoid mice eating cornmeal bedding and thus not being food motivated) and were acclimated to the new cage for one day before food deprivation. Food deprivation was conducted by removing food at 6:00 pm and returning it the next day after the animals completed the task. Acclimation to food deprivation was repeated for 2 days prior to the shaping phase of the study. Behavior tests were performed for each animal beginning at approximately 8:30 am each day in a skilled forelimb test apparatus with an adjustable vertical pellet holder (Maze Engineers) (*95*). The apparatus was 23-cm long, 10-cm wide, and 20-cm tall with a 0.5-cm wide vertical slit for the animals to reach through. Testing all 24 mice required about six hours. To reduce the effects of prolonged fasting on the animals' behavior, mice were randomized into three groups of eight and the testing order was rotated each day between groups. Mice are weighed immediately prior to behavioral testing to track weight loss.

In order to determine the number of animals necessary for this study we conducted a pilot study with three animals where we ran the mice through both the shaping and four days of training before and after laser ablation or stroke (n=2/group). Based on the effect size and variability in success observed in this pilot study, power analysis, run in GPower, indicated we needed 6 mice per group to determine a statistical difference at the $p = 0.05$ level. Because we knew that we would lose some animals due to surgical error and unwillingness to perform the task we increased our groups to 8 mice.

#### 4.2. Pellet Reaching Test

Shaping: Animals were placed in the behavior chamber for 20 minutes on 3 consecutive days. On day 1, sucrose pellets (20 mg, ScottPharma) were scattered within the chamber and placed in a tray outside the slit, within reach of the animals' paws but not their tongues to encourage grasp attempts. On days 2 and 3 of shaping, the pellets were only outside of the chamber for the mice to reach. Videos of the animals were manually scored by counting how many grasps were attempted by each paw to determine the animals' left/right paw preference.

Training: The animals were trained on their preferred paw for 8 days. Mice were placed in the behavior chamber for 20 minutes or until they reached 30 successful grasps. Sugar pellets were placed on the pellet holder 6 mm from the chamber wall and 3 mm lateral from the midline of the slit, contralateral to the animal's dominant paw. A blinded researcher, independent of those running the behavioral tests, manually scored videos of the animals' performance and tracked the total number of successful grasps, successful first attempt grasps, misses, and failed attempts where the pellet was knocked out of reach.

Testing of pellet reaching success occurred in three phases: before surgical intervention, in the week after surgical intervention, and three weeks after surgical intervention. Performance in the first phase guided which surgical intervention the animals should receive. Mice were removed from the study if they failed to reach a success rate of ~20% following initial training (2 mice). The remaining mice were divided into 3 groups to receive a craniotomy, a craniotomy with a regular ablation, or a craniotomy with a "severing ablation." Mice were allocated such that the average success rate, ratio of male to female mice, and ratio of mice from each original group of eight were approximately equal. Mice that received only a craniotomy were given a focal ischemic stroke after their three-week post-operative testing and the one week and three-week post-operative testing was repeated, so these same mice served as both positive and negative controls. Training was paused following the initial training phase for about a month and was resumed two days prior to any surgical procedures to refamiliarize the animals to the task. Performance remained high over this one month pause, consistent with previous findings (*96*).

#### 4.3. Surgery

Craniotomies for the behavior task differed from what was previously described only in location. Craniotomies were performed over the caudal forelimb area (CFA), contralateral to the animal's preferred paw, were centered ~1.5 mm lateral to bregma, and were 3 mm in diameter (*52, 95*) (Fig. S9A). For performing the ablation and severing ablation, the imaged field of view was centered ~1.5 mm from the sagittal sinus to cut a 1 mm diameter circle within the CFA. The regular ablation was created by cutting the same 1-mm diameter open cylinder spanning 70 to 550 μm depth, as described above (Fig. S9B). The severing ablation added a fully ablated 1-mm diameter plane at a depth of 500 μm below the cortical surface to this cylinder (Fig. S9C). The ablated layer was created by passing the beam path in a bidirectional manner from one end of the circle to another, with parallel lines spaced 25 μm apart.

Mice were given two days after surgery to recover and were given wet food during recovery. The pellet reaching task resumed for the second testing phase on the second full day post-operation although the animals were not food-restricted overnight. For day 3 post-operation onwards, food restriction resumed prior to behavioral tasks.

#### 4.4. Stroke

Mice that received a craniotomy only for the pellet reaching task had a subsequent stroke induced over the CFA after completing the three-week post-operative testing. Mice were given 50 μL retroorbital injection of Rose-Bengal (10 mg/mL) in saline. Under 2PEF imaging, a region was located within the CFA. A green CW laser at 530 nm was focused through the 0.95 NA 20x objective for 5 min to occlude vessels in the cortex (Supplementary Fig. 9D). The occluded region was confirmed using MESI. Mice were given the same 2-day rest period as was done for the first set of surgeries before resuming behavioral testing.

### 5. *Histological analysis of laser cuts to determine the chronic damage associated with ablation*

#### 5.1. Histology of laser cuts

Mice (n=12, 7 males and 5 females) from both the chronic imaging and behavioral study were perfused between 3-5 weeks after laser ablation using 4% PFA in PBS and brains were extracted from the skull before being stored in 4% PFA for 24 hours. Brains were then switched to a solution of 30% sucrose in PBS for another 24 hours. Brains were then cut into ~3-mm thick coronal slabs centered on the laser ablated region (marked on the dorsal surface with surgical ink). Tissues were washed in tap water and then processed on a Sakura Tissue-Tek VIP 6 tissue processor overnight where the tissues were infiltrated with a series of graded alcohol (70%, 80%, 95%, 100%), xylene, and finally melted infiltrating paraffin. Once processed, tissues were placed into a metal embedding mold filled with melted embedding paraffin, and fixed into the appropriate orientation by placing the mold on a cold plate to semi-solidify the paraffin. The cassette was then placed on top of the mold and filled with embedding paraffin, and the completed block was placed on the cold plate to solidify. The tissue was cut in 5-μm thick coronal sections which were mounted and stained using histochemical and immunohistochemical stains. Every other section was stained with hematoxylin and eosin. The intervening sections were stained for one of the following: Iba1 (microglia; Wako, Catalog #019-19741, 1:3000 dilution), Prussian blue (hemosiderin), luxol fast blue (myelin), olig2 (oligodendrocytes; ABCAM, catalog# ab109186, clone EPR2673,1:2000 dilution), and glial fibrillar acidic protein (astrocytes; Dako/Agilent, Catalog #Z0334, 1:3000 dilution). Each slide was examined by a board-certified veterinary pathologist, comparing the cut region with the contralateral cortex as a control.

### 6. *Data visualization and statistics*

Plots and graphs were created using Prism 8 (GraphPad) and MATLAB. For the correlation coefficient data (Fig. 1, Supplementary Figs. 3 and 4), correlation coefficients were calculated using MATLAB to compare the distant electrodes to the seizure focus, and violin plots comparing correlation coefficients were made in Prism 8 with the maximal width being at the highest density of values. Statistical significance for differences in the distribution of correlation coefficients between groups was determined using the Kolmogorov-Smirnov test. The 2-D contour plots were created using custom software in MATLAB and clustering used the Mahalanobis distance clustering method (*46*). Contours were created using a linear scale, distributing 30 contours between the maximum and minimum values of the data point density. All bar graphs (Figs. 1, 2, and 5, Supplementary Figs. 3 and 4) were created with the bar indicating the mean and whiskers showing standard deviation. Box and whisker plots (Figs. 1 and 2, Supplementary Figs. 6 and 7) were made with the box to include values between the 25$^{th}$ and 75$^{th}$ percentile with the whiskers extending to 1.5 times the interquartile range outside the box. Means were calculated using all values and are highlighted. Figure legends indicate when animals with outlier behavior were singled out with distinct color datapoints and means calculated with them excluded. Line graphs with shading that indicates the standard deviation were made in MATLAB (Figs. 2 and 4). Imaging data was analyzed in ImageJ and MATLAB.

Statistical analyses were done on Prism 8 and JMP Pro 15. When comparing groups, significance was determined using a one-way ANOVA with Tukey honest significant difference post hoc multiple comparisons correction for more than two groups and using an unpaired t-test for two groups. The number of epileptiform events that propagated, were attenuated, or were non-propagated were compared between control and laser cut mice using a Chi-squared test of association. To determine if the percentage of seizures that propagated changed over time, we ran a linear mixed effects model with a fixed effect of week and a random intercept and slope at the animal level for laser cut animals. For controls we ran a similar linear mixed model with the random intercept at the animal level. Fixed effect of week was tested using an F-test to determine if the trend over time was statistically significantly different than a slope of zero. For all linear and linear mixed effects models the residuals were assessed for normality and homogeneous variance; response variables were log transformed when needed. P-values were considered statistically significant when they were below 0.05. Statistical indicators are used consistently in figures, as follows: *$p<0.05$, **$p<0.01$, ***$p<0.001$, ****$p<0.0001$. Further details of each plot including sex differences, groups compared, statistical test, p-value, and explanatory notes are included in the figure legends.

### Supplementary Figures:

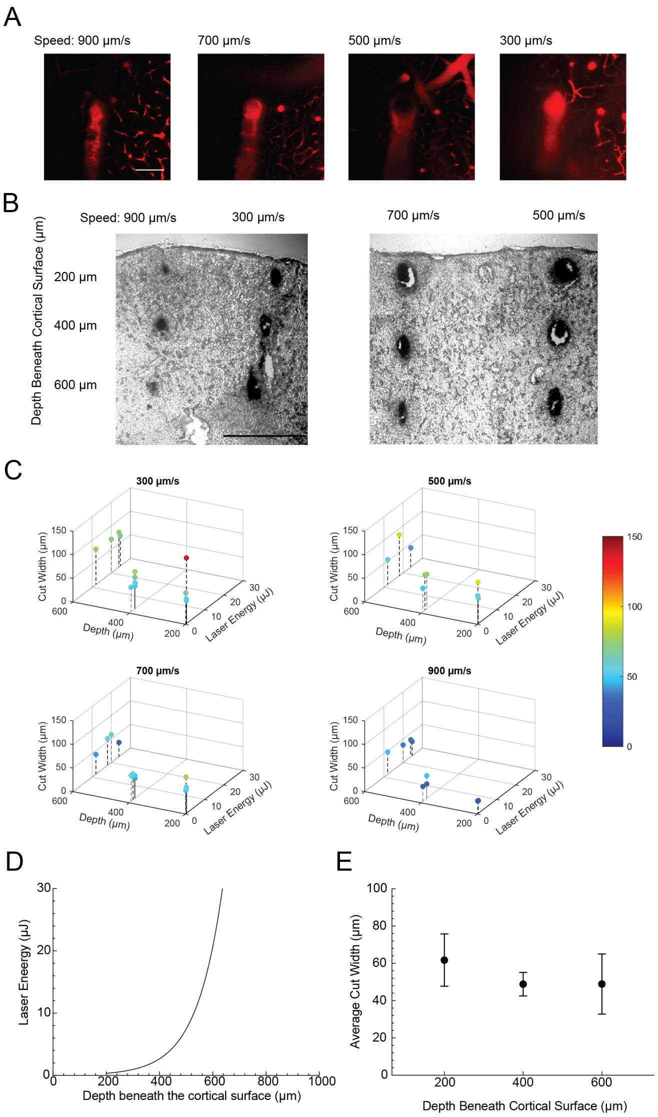


**Figure S1. Laser and cutting parameters were optimized to produce cuts that were uniformly ~50-µm wide as a function of depth into the cortex. (A)** Two-photon images taken immediately after ablation at 600 µm below the cortical surface for cuts produced with 12 µJ and at several translation speeds (scale bar: 100 µm). Texas red dextran labeled the vasculature, and the extravasation of blood plasma into the brain tissue at the cut location is apparent. **(B)** Light microscope images of 3,3′-diaminobenzidine (DAB) stained coronal slices at the location of laser cuts produced at different speeds across several depths, with the laser energy varying with depth as shown in panel D. Cuts in each column were made with at the same speed, as indicated above the column (scale bar: 100 µm). **(C)** 3D plot of cut width as a function of depth and laser energy. Colors are a secondary indicator of cut width to make the results more apparent. Above each plot is the speed with which the stage was moved. **(D)** Laser energy used as a function of depth into the cortex, showing the exponential increase in energy needed to overcome optical scattering losses (based on results in Nguyen, et al. (*42*). While this prior work established appropriate laser energy as a function of depth, optimal laser translation speeds through the tissue that were as fast as possible while producing uniform cuts were not determined. **(E)** Cut width as a function of depth beneath cortical surface for 700 µm/s translation speed while using the depth-dependent energy shown in panel D. Values at each depth are an average of the cut widths for the energies that were used and fit the curve in part D (average: 55 µm).

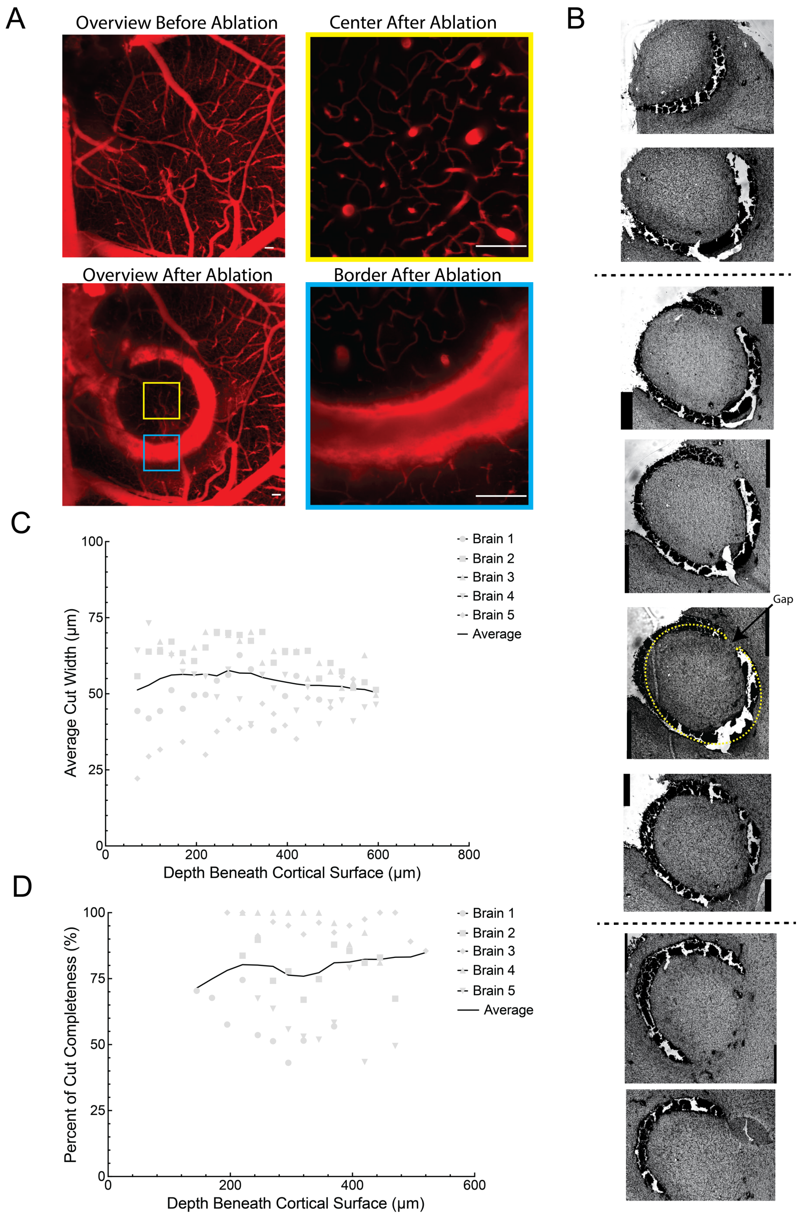


**Figure S2. Cuts at depths ranging from 550 µm to 70 µm below the cortical surface were ~85% complete with a ~55 µm cut width. (A)** Two-photon images taken before and after laser ablation with blood plasma labeled by Texas Red-dextran. Top left image: before ablation; Bottom left image: after ablation; Right images from the center (top) and edge (bottom) of the cut at higher magnification after ablation. Scale bars: 100 µm. **(B)** Images of DAB-stained horizontal brain slices through the circular ablated region. Images between 170 and 450 µm (indicated by the black dotted lines) were used to measure cut completeness. Slices from above and below this region often contained tissue above or below where we cut, due to a small tilt of the cutting plane relative to the brain surface, and were ignored. The completeness of the cut (e.g. yellow dotted line in the middle picture) and gaps in the cut (indicated by an arrow in the middle picture) were identified. **(C)** Average cut width and **(D)** cut completeness as a function of depth below the cortical surface, showing data from five separate brains in grey and smoothed average lines in black.

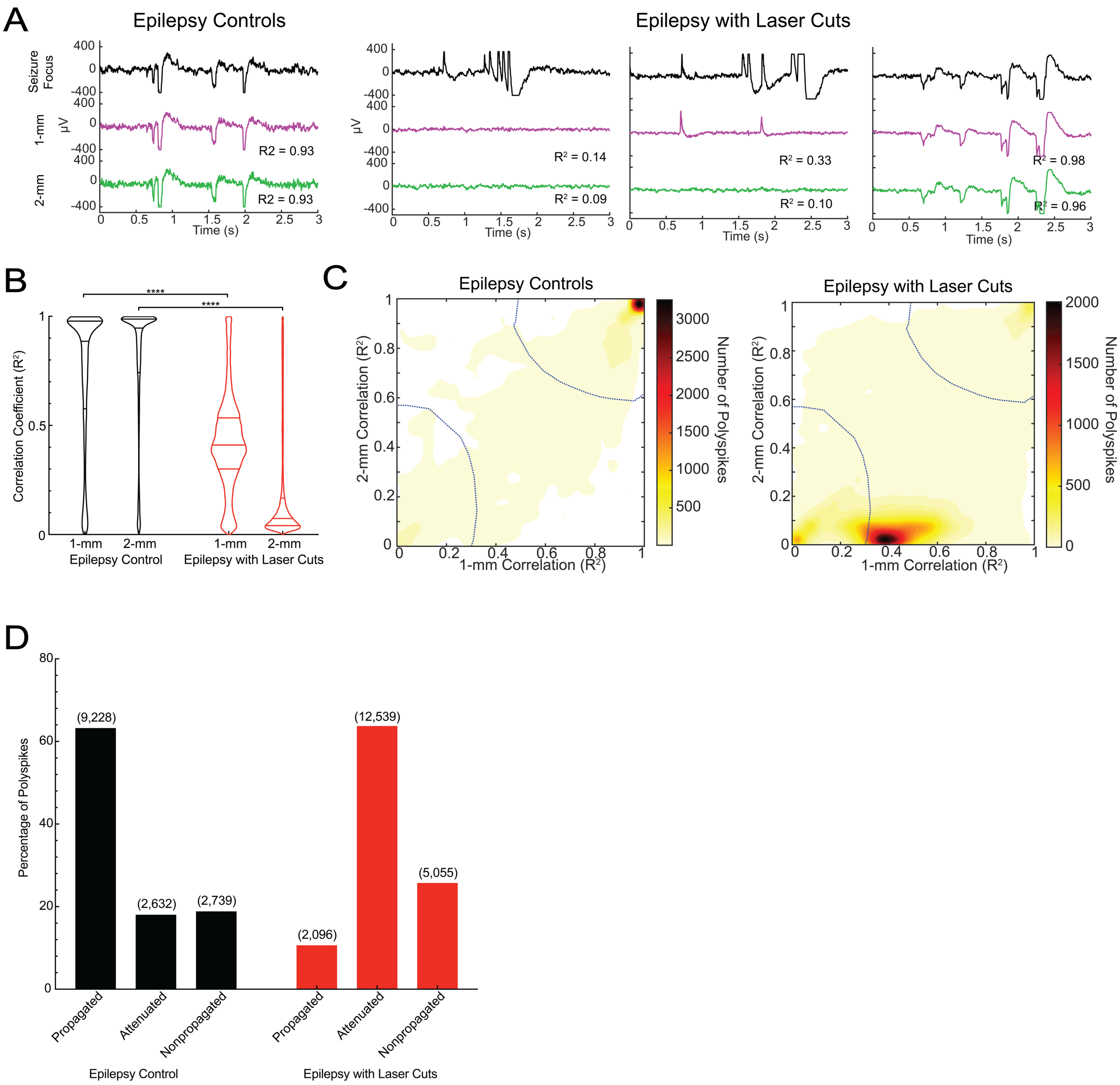


**Figure S3. Laser cuts significantly blocked polyspike propagation. (A)** Examples of polyspikes from epilepsy controls (left: propagated) and epilepsy with laser cuts (right: non-propagated, attenuated, and propagated, respectively from left to right), with the corresponding correlation coefficients indicated below each trace. **(B)** Correlation coefficients comparing the events at 1-mm and 2-mm electrodes with the seizure focus between controls and laser cut animals (****$p<0.0001$ Kolmogorov-Smirnov test). **(C)** 2-D contour plot of epilepsy controls (left) and epilepsy with laser cuts (right) with clusters bounded by blue lines: propagated (top right), attenuated (middle), non-propagated (bottom left). **(D)** Bar graph representing the percent of total polyspikes in each of the clusters for epilepsy control and epilepsy with laser cuts (red) (numbers above columns are the number of polyspikes in each of the clusters). Clustering was different between laser cut and control groups ($P<0.0001$, Chi-squared test of associations).

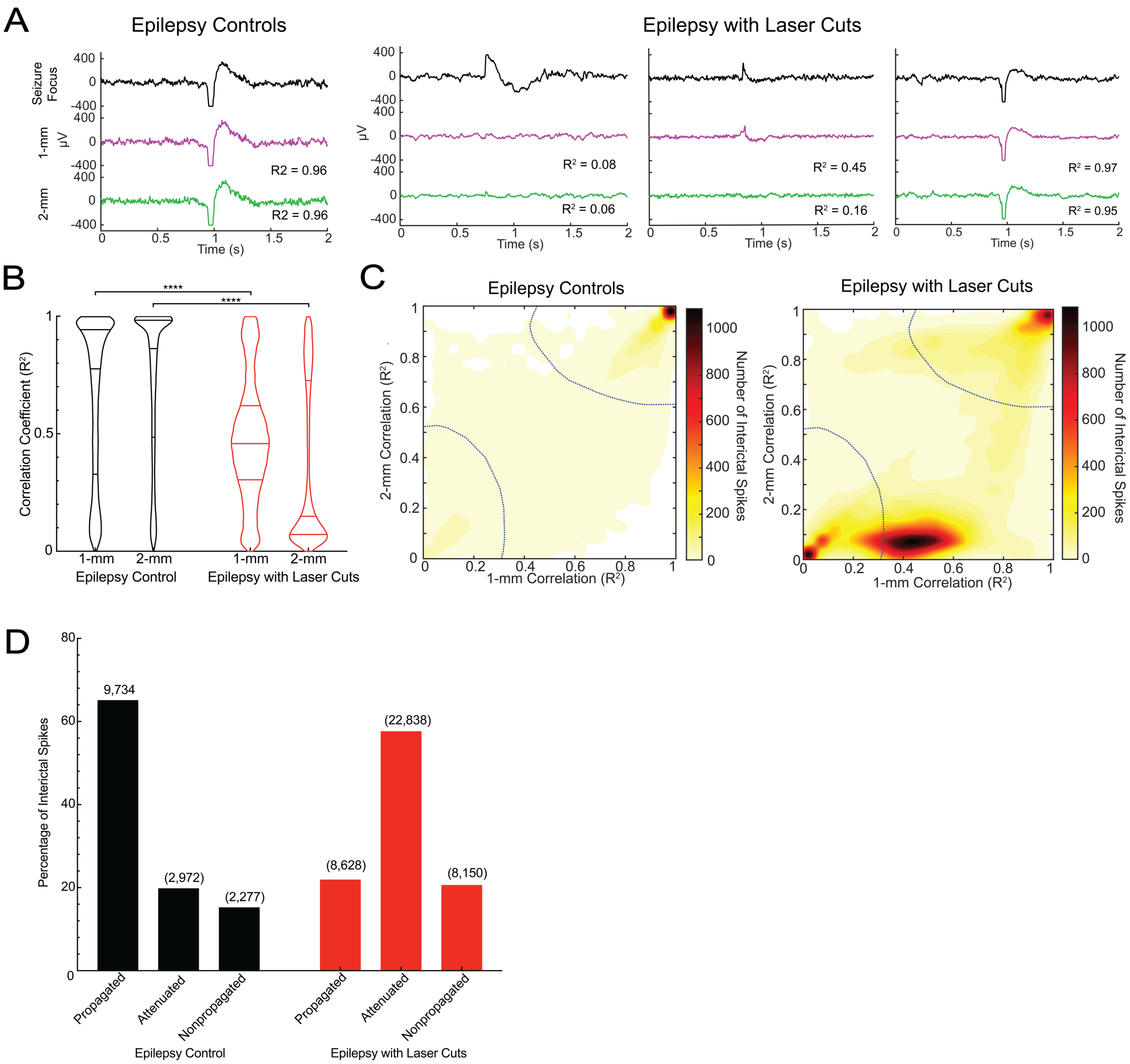


**Figure S4. Interictal spike propagation was reduced by laser ablation surrounding a seizure focus. (A)** Examples of interictal spikes from epilepsy controls (left: propagated) and epilepsy with laser cuts (right: non-propagated, attenuated, and propagated, respectively from left to right), with the corresponding correlation coefficients indicated below each trace. **(B)** Correlation coefficients comparing the events at 1-mm and 2-mm electrodes with the seizure focus between controls and laser cut animals (****p<0.0001 Kolmogorov-Smirnov test). **(C)** 2-D contour plot of epilepsy controls (left) and epilepsy with laser cuts (right) with clusters bounded by blue lines: propagated (top right), attenuated (middle), non-propagated (bottom left). **(D)** Bar graph representing the percent of total interictal spikes in each of the clusters for epilepsy control and epilepsy with laser cuts (red) (numbers above columns are the number of interictal spikes in each of the clusters). Clustering was different between laser cut and control groups (P<0.0001, Chi-squared test of associations).

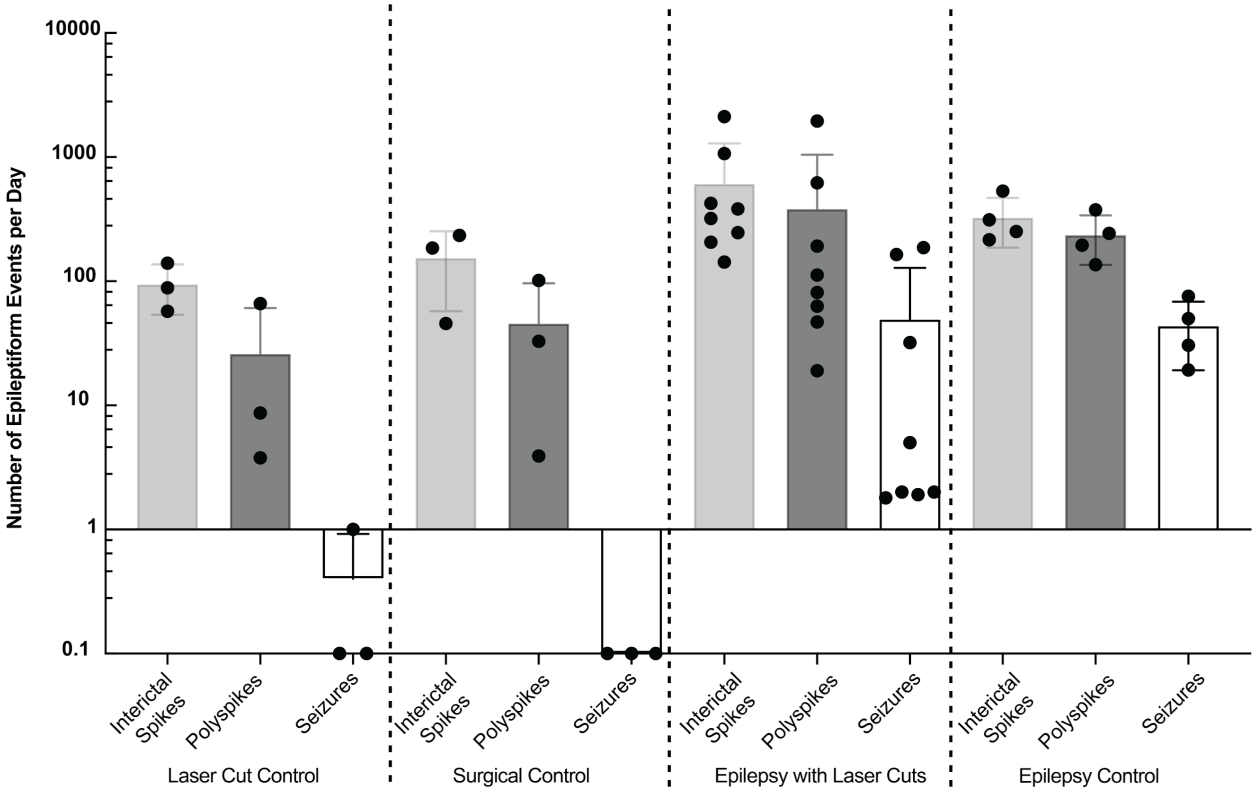


**Figure S5. Incidence of epileptiform events across all experimental groups show that iron chloride was the inciting factor for seizure induction, and laser cuts did not cause seizures.** Comparison of epileptiform events across laser cut control, surgical control, epilepsy with laser cuts, and epilepsy control. Animals with zero recorded seizures were set to 0.1 events/day to facilitate visualization on a log plot.

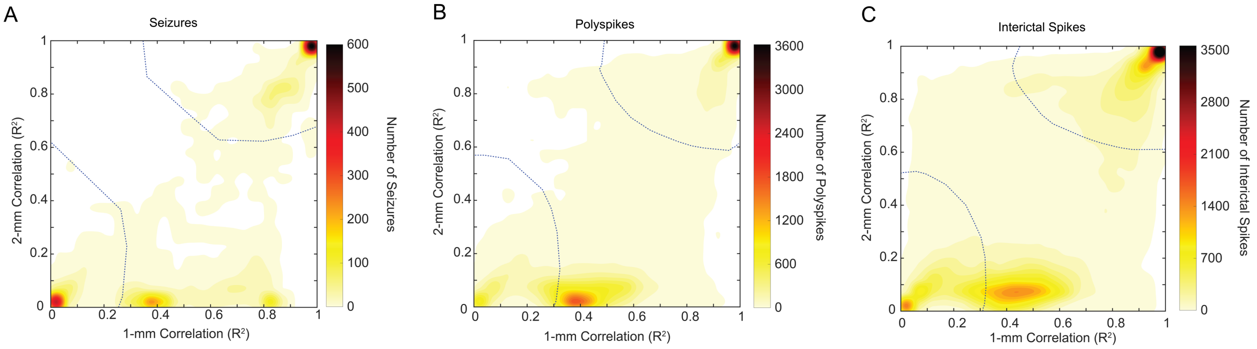


**Figure S6. Combination of all epileptiform events across epilepsy control and epilepsy with laser cut animals indicate three clear clusters of events: propagated, attenuated, and non-propagated. (A-C)** 2-D contour map of correlation coefficients combining both epilepsy control and epilepsy with laser cuts for (A) seizures, (B) polyspikes, and (C) interictal spikes.

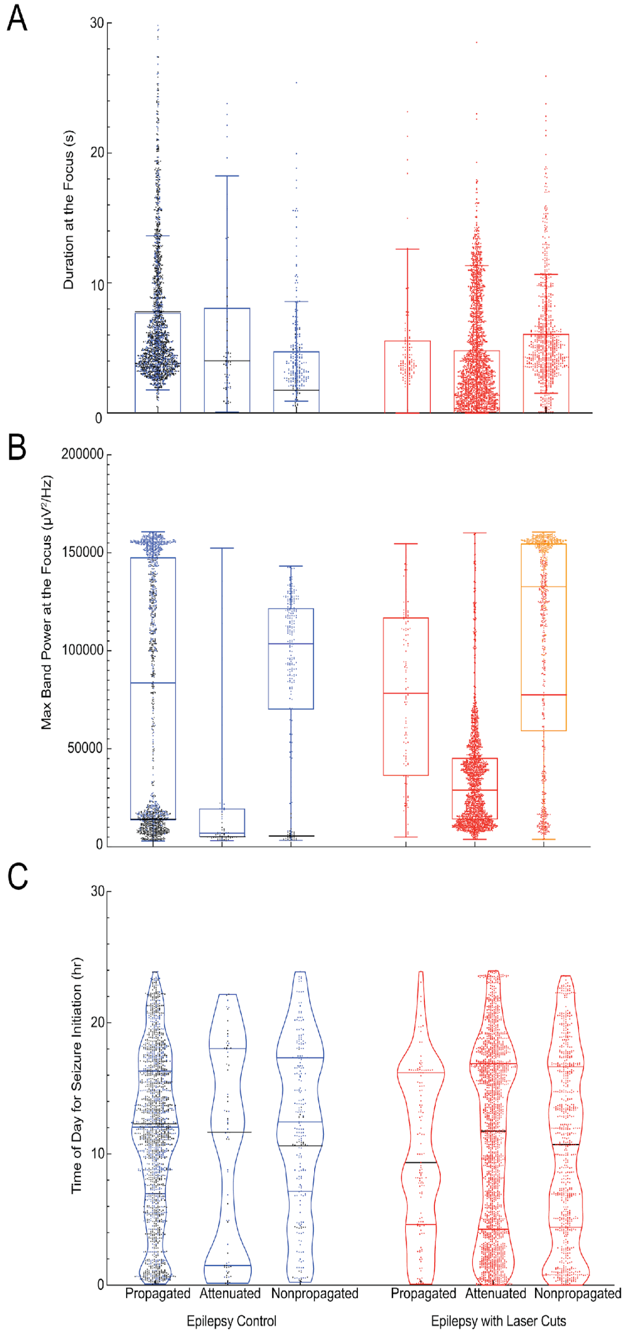


**Figure S7. Seizure metrics at the focus suggest that non-propagated seizures in epilepsy with laser cut animals would have propagated without laser cuts. (A)** Seizure duration at the focus compared across clusters for epilepsy control and epilepsy with laser cut groups. **(B)** Max band power at the seizure focus compared between epilepsy control and epilepsy with laser cuts (red) over the three clusters. **(C)** Time of day that seizures occurred across groups and clusters. Data points from outlier mice are shown with different colors to avoid hiding average trends (blue: epilepsy control outlier – black mean indicators show trend; orange: epilepsy with laser cuts outlier – red mean indicators show trend).

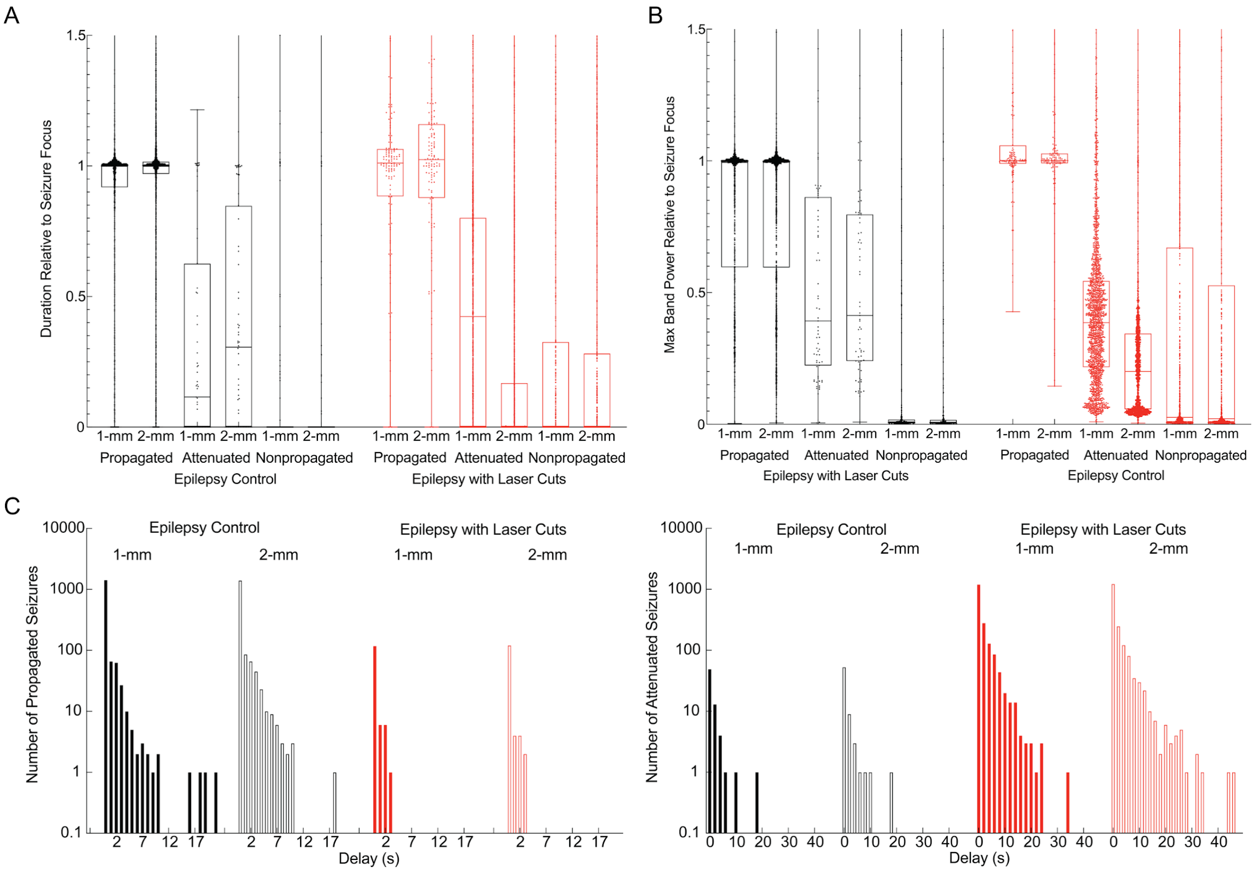


**Figure S8. Seizure metrics at distant electrodes show our method of clustering accurately portrays the physiological seizure propagation. (A)** Seizure duration and **(B)** Max band power at the distant electrodes, as a fraction of the value at the focus, for epilepsy control and epilepsy with laser cuts (red) mice and across propagated, attenuated, and non-propagated seizures. **(C)** Delay between seizure initiation at the focus and arrival at the distant electrodes for epilepsy control and epilepsy with laser cuts groups, illustrating propagated (left) and attenuated (right) seizures (filled bars – 1-mm, open bars – 2-mm).

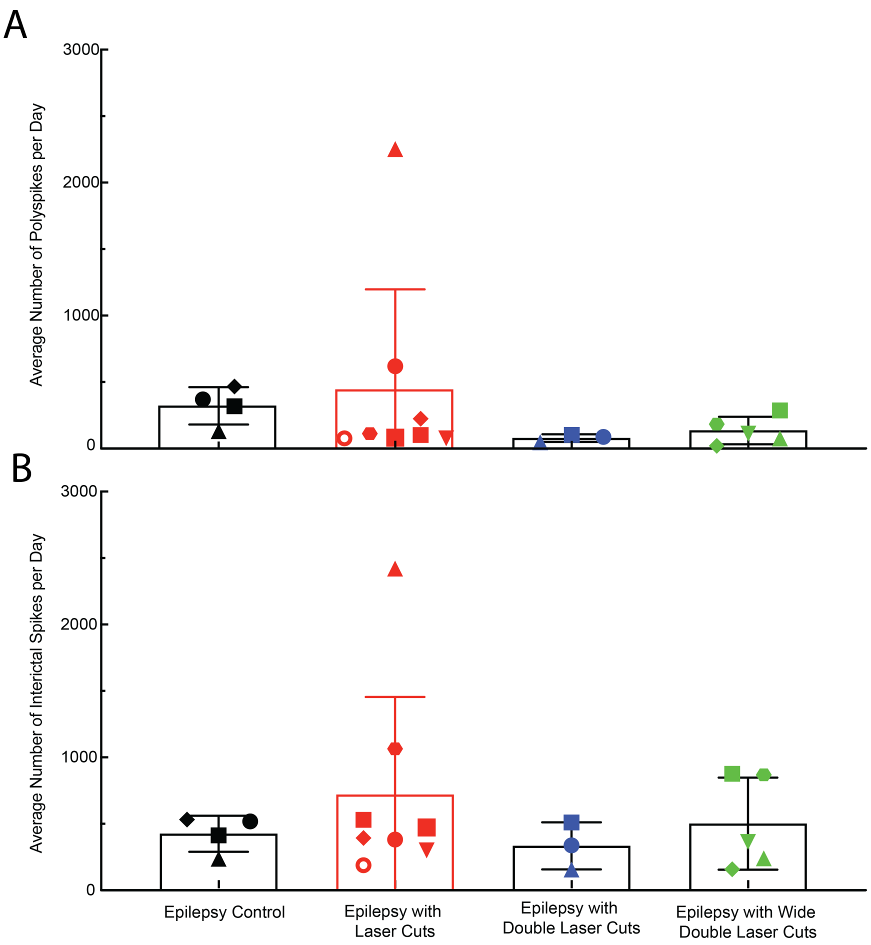


**Figure S9. Laser cuts did not reduce polyspike or interictal spike frequency.** Average number of **(A)** polyspikes and **(B)** interictal spikes per day for epilepsy control, epilepsy with laser cuts, epilepsy with double laser cuts, and epilepsy with wide double laser cuts groups.

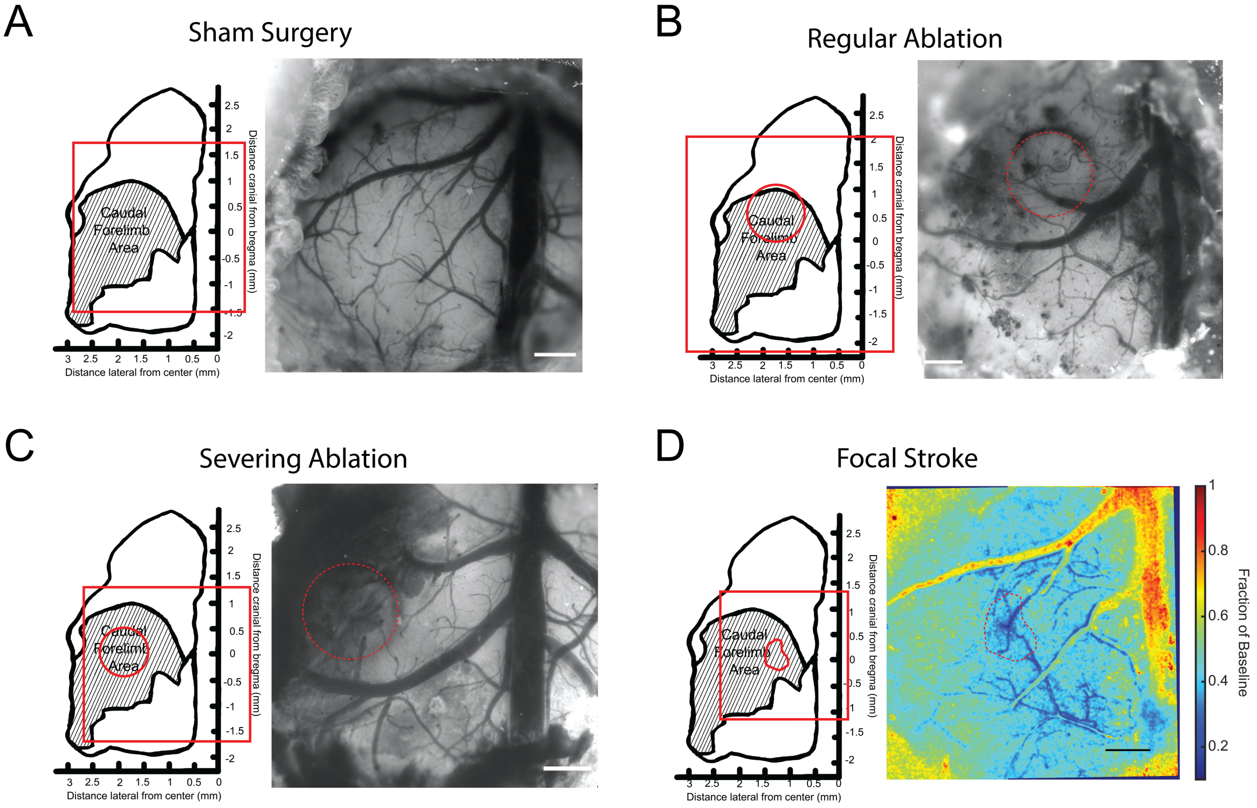


**Figure S10. Treatments used for behavioral testing were confirmed to be in the caudal forelimb area of motor cortex. (A-C)** Green light images showing blood vasculature on the right with a map indicating the craniotomy location with a red box and outlining the location of laser cuts in B & C with a red circle. **(D)** Focal stroke with the same map and indicators on the left and heat map based on laser speckle contrast imaging on the right. The stroked area is highlighted with a red dotted line and indicated by a lower fraction of baseline blood flow. Scale bars: 500 µm.

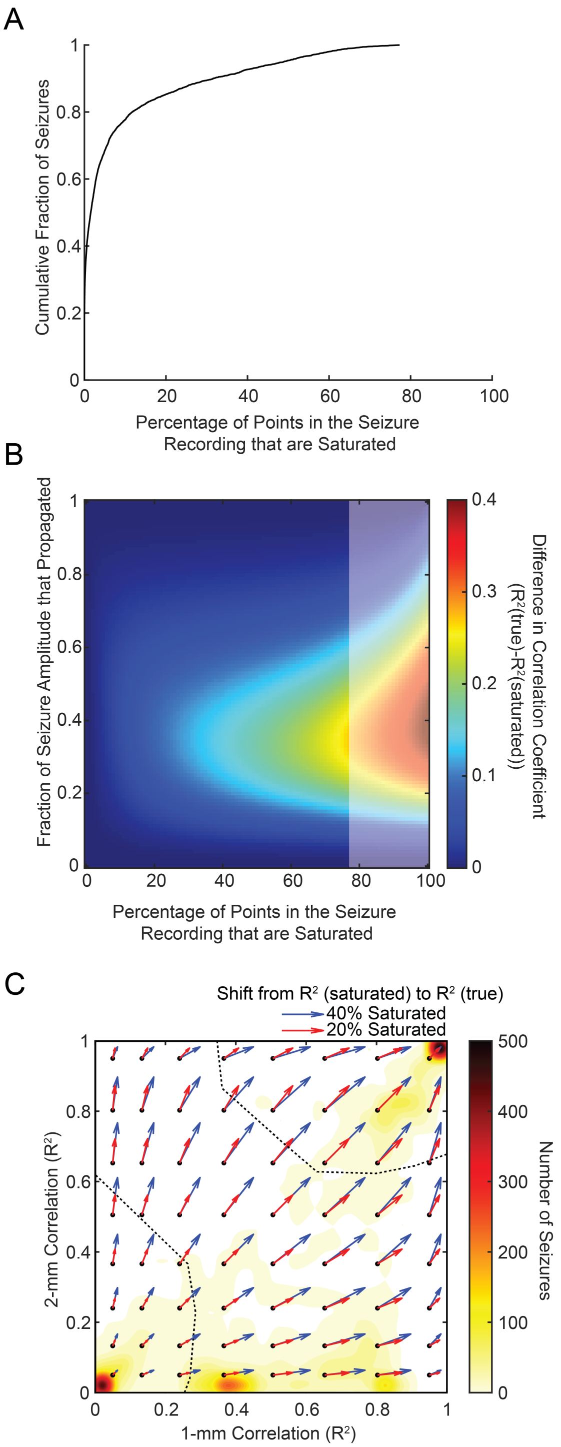


**Figure S11. Saturation of ECoG signal during epileptiform events decreases the correlation coefficient between the seizure focus and distant electrodes but does not significantly impact classifications. (A)** The cumulative distribution of all seizures that fall below a given percentage of samples within the signal that are saturated. **(B)** The theoretical differences of correlation coefficients at varying levels of saturation and seizure amplitude breakthrough. No experimental data is in the lightly shaded region. **(C)** A displacement map of the measured correlation coefficients (black dots) and the distances to the true measurements under a 20% (red) and 40% (blue) saturation condition overlaid onto the contour distribution of seizure correlation measurements.

**Supplementary Movies**

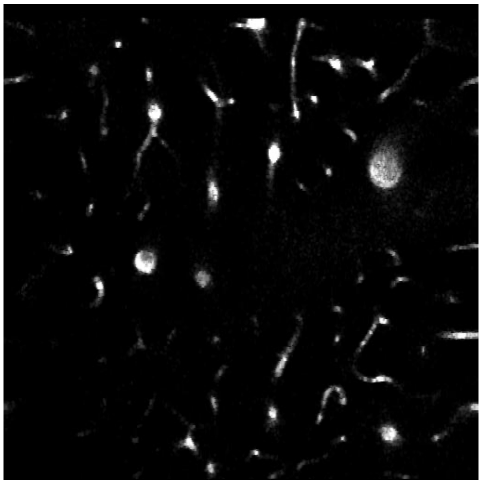

**Movie S1. Two-photon microscopy video of the production of a 1-mm dimeter cylindrical laser cut through cortical layers II-IV (550 µm to 70 µm).** Blood vessels are labeled with Texas Red dextran. Video starts at the surface of the brain and then steps down to 550 µm below the cortical surface where laser ablation begins. Ablation moves in a circular pattern, pauses to move dorsally 10 µm, and then repeats the circle again. This process is repeated from 550 µm to 70 µm below the cortical surface.

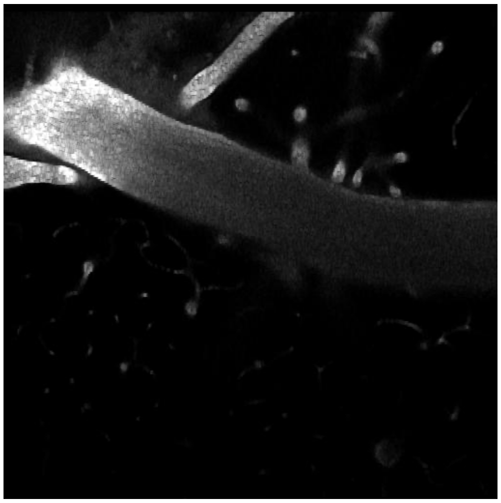

**Movie S2. Two-photon imaging depicting closely spaced double laser cuts (100-µm spacing) made from 550 µm to 70 µm below the cortical surface.** Video follows the same trajectory as Supplementary Video 1 but completes the inner circle (1-mm diameter) and then the outer circle (1.2-mm diameter) at every level before moving dorsally.

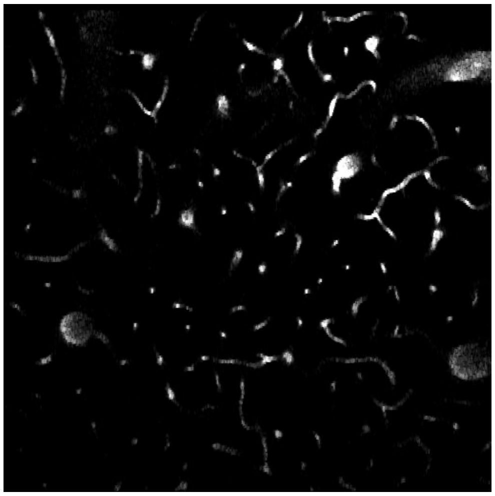

**Movie S3. Two-photon imaging illustrating widely spaced double laser cuts (300-µm spacing) made from 550 µm to 70 µm below the cortical surface.** Video follows the same trajectory as Supplementary Video 1 but completes the inner circle (1-mm diameter) and then the outer circle (1.6-mm diameter) at every level before moving dorsally.

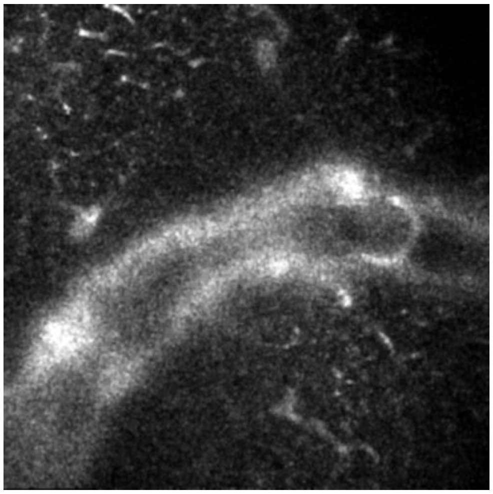

**Movie S4. Two-photon imaging demonstrating the severing of vertical connections for the severing ablation group for the pellet reaching task.** Blood vessels are labeled with Texas Red dextran. Single laser ablation from 550 µm to 70 µm below the cortical surface was completed as with above (Supplementary video 1). Severing of a single layer at 500 µm below cortical surface is depicted here by panning the laser back and forth with 25 µm between lines.
